# *Trypanosoma brucei* bloodstream form mitochondrion is capable of ATP production by substrate phoshorylation

**DOI:** 10.1101/2023.05.26.542429

**Authors:** Taleva Gergana, Husova Michaela, Panicucci Brian, Hierro-Yap Carolina, Pineda Erika, Marc Biran, Martin Moos, Petr Šimek, Bringaud Frederic, Zíková Alena

**Author notes:** these two authors contributed equally (Taleva Gergana, Husova Michaela).

## Abstract

The bloodstream form *Trypanosoma brucei* maintains essential mitochondrial membrane potential (ΔΨm) through the reverse activity of FoF1-ATP synthase. The ATP that drives this activity is thought to be generated by glycolysis and imported from the cytosol via an ATP/ADP carrier (AAC). We have shown that this carrier is the only carrier that can import ATP into the mitochondrial matrix to power the F_o_F_1_-ATPase. Contrary to expectations, its deletion has no effect on parasite growth, virulence and levels of ΔΨm, suggesting that ATP is produced intramitochondrially by substrate phosphorylation pathways. Therefore, we knocked out the succinyl-CoA synthetase (SCoAS) gene, a key enzyme that produces ATP through substrate phosphorylation. Its absence resulted in changes in the metabolic landscape of the parasite, lower virulence, and reduced mitochondrial ATP content. This minimal mitochondrial ATP pool was maintained by AAC activity as evidenced by the 25- fold increase in sensitivity of the mutant parasites to AAC inhibitor carboxyatractyloside. Under nutrient-limited conditions, suppression of SCoAS expression by RNA interference negatively affected cell growth and levels of ΔΨm. We concluded that the bloodstream mitochondrion is capable of generating ATP via substrate phosphorylation pathways, the importance of which depends on environmental conditions.

## Introduction

*Trypanosoma brucei*, the causative agent of African trypanosomiasis, is a master of metabolic adaptations to its environment. This unicellular parasite, which undergoes a complex digenetic life cycle in the insect and mammalian hosts, metabolically adapts to the host environment and the nutrients offered [1]. In terms of energy metabolism, the insect forms of the parasite consume mainly amino acids (e.g. proline, threonine), which are oxidized in its single mitochondrion to succinate, acetate and alanine [2-4] and generates ATP by both oxidative and substrate phosphorylation pathways [5-8]. On contrary, the bloodstream form (BSF) generates the majority of cellular ATP in the cytosol by glycolysis and excludes its mitochondrion as the powerhouse of the cell [9]. Therefore, the bloodstream form is a rare example of an aerobic organism not generating most of its ATP by oxidative phosphorylation. This is possible because this form resides in the glucose-rich environment of its mammalian host containing enough substrate for highly efficient glycolysis [10]. In addition, the BSF does not assemble the proton-pumping electron transport chain (ETC) complexes III and IV, but expresses the trypanosoma alternative oxidase (TAO or AOX) [11], an enzyme that takes over the role of these ETC complexes, but does not contribute to proton motive force. Therefore, the presence of AOX uncouples the parasite respiration from generation of mitochondrial membrane potential (ΔΨm) that powers oxidative phosphorylation by F_o_F_1_-ATP synthase, hence no ATP is produced by this pathway.

Why *T. brucei* BSF makes such an extreme metabolic switch from canonical ETC complexes III and IV to AOX is not clear, but since the complex III is one of the largest producers of ROS during aerobic metabolism, it is a possible adaptation to lower the amount of these potentially harmful molecules [12]. The consequence of this electron detour is that AOX cannot contribute to the ΔΨm, an essential attribute of a healthy mitochondrion that not only drives ATP production but is also critical for protein import into the organelle and for mitochondrial ion exchange. One would assume that in the absence of the proton-pumping complexes III and IV, ΔΨm would be maintained by the ETC complex I, another powerful proton pump fed by electrons from reduced mitochondrial NADH, but the genetically generated evidence does not support its role in generating proton motive force in the bloodstream form [13].

Uniquely, the BSF ΔΨm is maintained by the reverse activity of F_o_F_1_-ATP synthase, which hydrolyzes ATP and uses the released energy to pump protons across the inner mitochondrial membrane [14, 15]. The reverse activity of this complex is well known in the aerobic eukaryote world, but is only used for a short time to overcome rapid changes in the environment (e.g. sudden hypoxia or anoxia) involving impaired respiration and therefore mitochondrial membrane depolarization [16]. Under these conditions, first the F_o_F_1_-ATP synthase reverses and hydrolyzes ATP that is supplied by mitochondrial substrate phosphorylation, a rescue mechanism that protects against cytosolic ATP depletion [17, 18]. However, if ΔΨm is reduced even more and intramitrochondrial ATP/ADP ratio decreases, the ATP/ADP carrier (AAC) also reverses supplying now the F_o_F_1_-ATP synthase pump with the cytosolic ATP, a condition that can quickly precipitate to cell death if not stopped by an inhibitory peptide IF1 [19, 20].

A. *T. brucei* is therefore a rarity that exploits this reverse activity over the long term. The hydrolytic activity of F_o_F_1_-ATP synthase appears to be the only entity that generates ΔΨm in this parasite, as RNAi silencing of its subunits causes a decrease in ΔΨm within 24 hours [15, 21-23], and inhibition of this complex by the peptide inhibitor TbIF1 decreases ΔΨm below the viability threshold within 12 hours [24]. Thus, the bloodstream form F_o_F_1_-ATP synthase is not an ATP-producing enzyme but an ATP-consuming enzyme. The question is through which metabolic pathways this molecular nanomachine is supplied with ATP. There are at least two possibilities: either ATP is taken from the cytosol and imported into the mitochondrial matrix by (an) ATP/ADP carrier(s) [25, 26], or the mitochondrion produces the ATP itself through substrate phosphorylation pathways. Because the mitochondrion of the bloodstream form is metabolically poor when compared with the insect forms [27], it was assumed that the organelle does not participate in the ATP production and that the glycolytically produced ATP is imported from the cytosol. However, there is no direct experimental evidence for this assumption.

In addition, recent metabolomic and proteomic data suggest that the metabolic potential of the BSF parasite mitochondrion may be greater than originally thought [28-30]. For example, a portion of glucose-derived pyruvate and threonine are further metabolized to acetate, an essential precursor for de novo fatty acid synthesis [31]. Glucose-derived pyruvate and threonine are metabolized by pyruvate-and threonine dehydrogenases (PDH and TDH), respectively, leading to the formation of acetyl-CoA. This energy-rich compound is rapidly converted to acetate by two redundant pathways. The first utilizes acetyl-CoA thioesterase (ACH), and the second utilizes acetate:succinate-CoA transferase (ASCT), which is coupled to succinyl-CoA synthetase (SCoAS) activity and simultaneously produces mitochondrial ATP [32] (Figure 1). Isotope-labeled metabolomic data have also shown production of succinate that is not derived from glucose, suggesting metabolism of other carbon sources, such as amino acids [30]. Interestingly, the bloodstream form consumes much glutamine from the medium [29]. Glutamine-derived α- ketoglutarate can be converted by α-ketoglutarate dehydrogenase to succinyl-CoA, which is the substrate for ATP-producing SCoAS. Moreover, α-ketoglutarate can be produced by amino acid transaminases. Thus, it appears that the bloodstream form mitochondrion may be capable of intramitochondrial ATP production (Figure 1).

**Figure 1.**
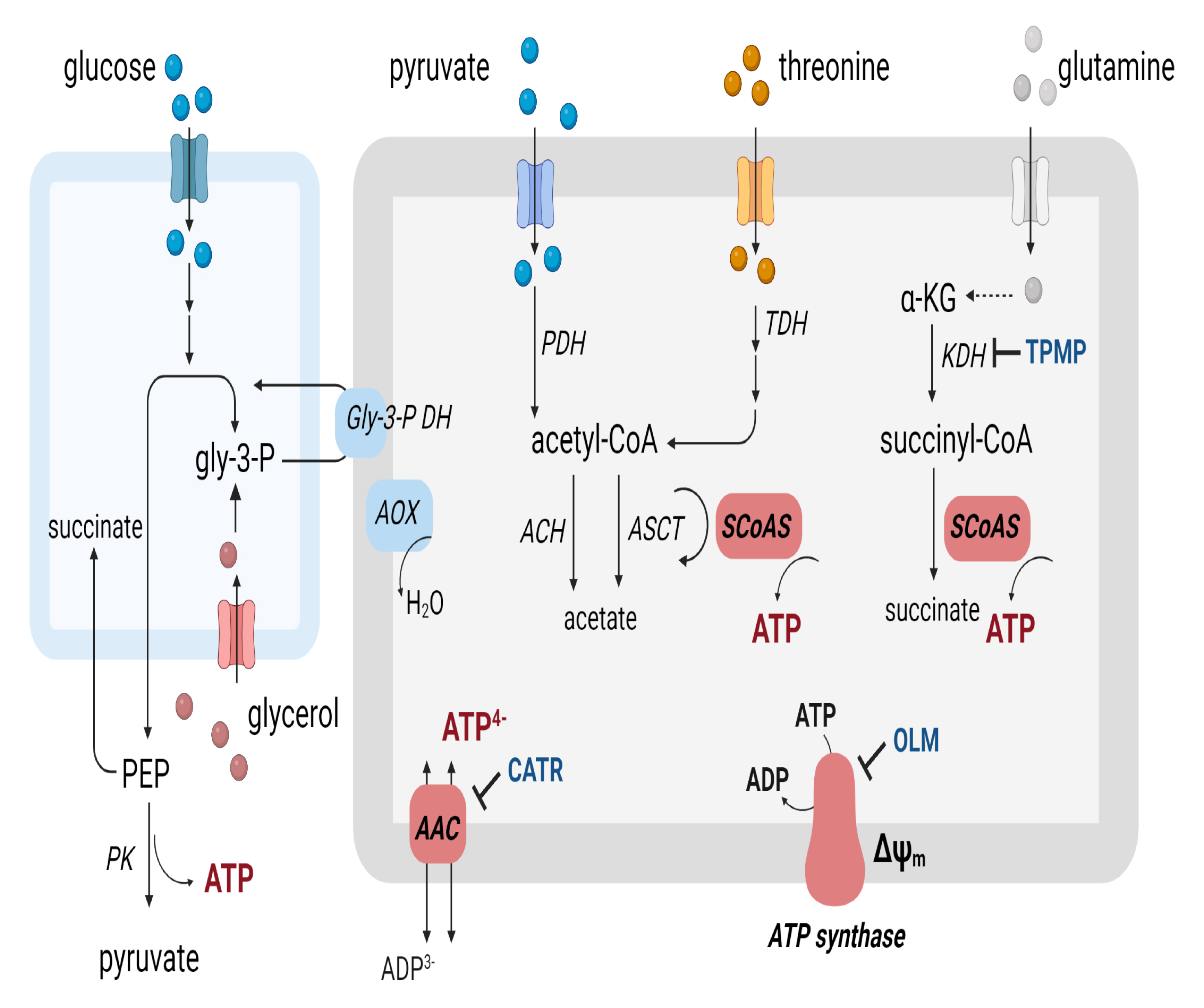
Schematic simplified representation of possible metabolic pathways linked to cytosolic and mitochondrial ATP production. Enzymes are: ACH, acetyl-CoA thioesterase, AOX, alternative oxidase, ASCT, acetate:succinate CoA-transferase, Gly-3-P DH, glycerol-3- phosphate dehydrogenase, KDH, α-ketoglutarate dehydrogenase, PK, pyruvate kinase, PDH, pyruvate dehydrogenase, SCoAS, succinyl-CoA synthetase, TDH, threonine dehydrogenase. Abbreviations: α-KG, α-ketoglutarate, ΔΨm, mitochondrial membrane potential, AAC, ATP/ADP carrier, CATR, carboxyatractyloside, OLM, oligomycin, TPMP, methyltriphenylphosphonium.

Since sequestration of cationic drugs (e.g. isometamidium and diminazene) commonly used to treat animal African trypanosomiases depends on the high levels of ΔΨm [33, 34] and resistance to some of these drugs is linked to reduced ΔΨm [35-37], it is crucial to understand the molecular mechanisms responsible for maintenance of the ΔΨm in trypanosomes. To determine which molecular entity (-ies) supplies (-y) ATP to the reversed F_o_F_1_-ATP synthase, we generated null mutants for the ATP/ADP carrier (AAC) and also for the SCoAS, an enzyme responsible for mitochondrial substrate phosphorylation, and tested the effects of their absence on viability and the mitochondrial metabolism and bioenergetics of *T. brucei* bloodstream form.

## Results

### ATP/ADP carrier is dispensable in BSF *T. brucei in vitro* and *in vivo*

The ATP/ADP carrier (AAC, originally named MCP5 [25]) of *T. brucei* is represented by three identical and consecutive genes (Tb927.10.14820, −14830, −14840) in the parasite genome. To test whether cell viability of BSF depends on the presence of AAC, we removed all three genes by homology recombination, resulting in an AAC double knock-out mutant (AAC DKO) (Figure 2A). We verified the correct genomic integration of the two cassettes containing antibiotic resistance genes by PCR (Figure 2B) and by Western blot using a specific polyclonal antibody against *T. brucei* AAC [26] (Figure 2C). The AAC DKO mutants showed no significant effect on growth when grown in commonly used HMI-11 medium containing a high concentration of glucose (25 mM) (Figure 2D), nor in the simplified Creek minimal medium (CMM) containing glucose at a lower concentration (10 mM) and therefore resembling better the in vivo environment of BSF [38](Figure 2E). We also tested whether the mutant parasites were virulent when introduced into an animal model. We infected two groups of BALB/c mice and monitored survival and parasitemia over several days with the parental (BSF427) and AAC DKO parasites. Neither group of infected mice survived beyond day 6, indicating that the AAC DKO mutants are fully virulent in the mouse model (Figure 2F). These results indicate that AAC is dispensable for BSF cell viability.

**Figure 2.**
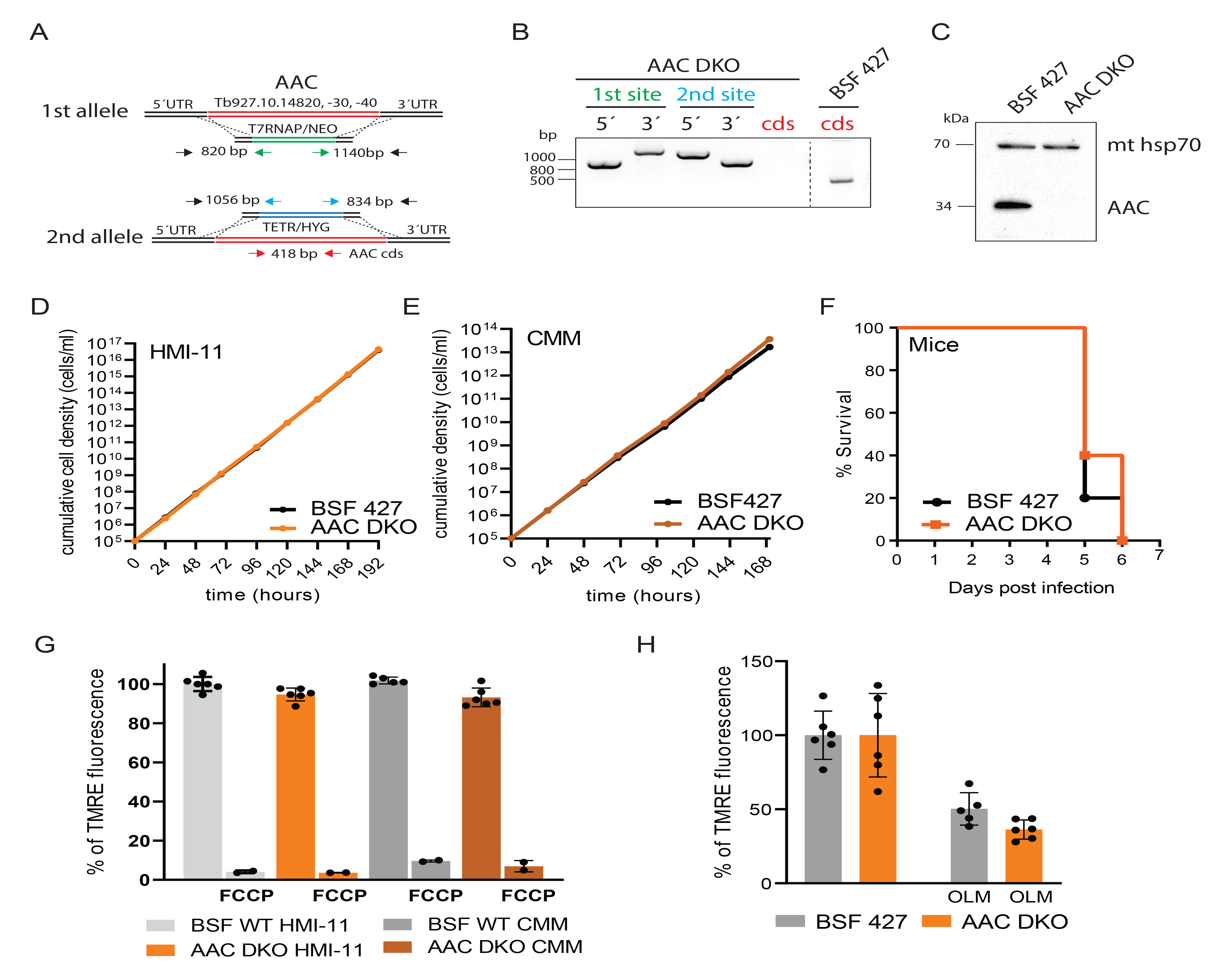
The ATP/ADP carrier is dispensable for BSF *T. brucei* viability and for maintaining the ΔΨm. A. The strategy to generate AAC DKO involved replacement of both alleles with T7 RNA polymerase and tetracycline repressor linked to resistance genes conferring neomycin and hygromycin resistance, respectively. B. PCR verification for the elimination of all AAC alleles in AAC DKO cell line. The primers used are color-coded in A. C. Immunoblot analysis of AAC DKO cells using specific anti-AAC antibody. Immunodetection of mitochondrial hsp 70 served as a loading control. D. Growth of AAC DKO cells compared to wild-type BSF 427 in HMI-11 measured for 8 days. E. Growth of AAC DKO cells compared to wild-type BSF 427 in CMM medium measured for 7 days. F. The survival rate of 5 female BALB/c mice which were intraperitoneally infected with AAC DKO and wild-type BSF 427 parasites. The infected mice were monitored for 6 days. G. Flow cytometry analysis of TMRE-stained AAC DKO and BSF 427 cells grown in HMI- 11 or CMM medium to measure ΔΨm. The addition of FCCP served as a control for ΔΨm depolarization (+FCCP). (means ± s.d., n= 6) H. Flow cytometry analysis of TMRE-stained AAC DKO and BSF 427 cells grown in HMI- 11 medium and treated with 250 ng/ml of oligomycin (+OLM) for 24 hours before the analysis. (means ± s.d., n= 6)

The predicted role of AAC in BSF parasites is to function in reverse, supplying the mitochondrial matrix with cytosolic ATP. This molecule is then hydrolyzed by F_o_F_1_-ATP synthase to maintain the essential ΔΨm. We investigated whether the absence of AAC affects ΔΨm in living cells using a fluorescent lipophilic dye, tetramethylrhodamine ethyl ester (TMRE) in non-quenching mode by flow cytometry. We found no difference in fluorescence intensity averaged over the entire cell population of BSF427 and AAC DKO cells grown in either HMI-11 or CMM demonstrating that AAC DKO maintains the ΔΨm at the same level as BSF 427. The treatment with FCCP, a protonophore, induced membrane depolarization as expected (Figure 2G). To test if the AAC DKO maintains the ΔΨm by the reversed activity of F_o_F_1_-ATP synthase, the BSF427 and AAC DKO cells were incubated for 24 h with a sublethal concentration of oligomycin, the F_o_F_1_-ATP synthase inhibitor (250 ng/ml, ∼0.5 of the EC_50_ for BSF427 [21, 39]). This treatment did not affect the doubling time of BSF427 and AAC DKO (BSF 427: 6 ± 0.2 hours, AAC DKO: 6.3 ± 0.3), and resulted in a similar reduction of ΔΨm in BSF427 and AAC DKO mutant in values reaching 50±11% and 64±7%, respectively (Figure 2H). Moreover, the AAC DKO mutant remained sensitive to the treatment with oligomycin, as tested by the Alamar Blue assay, with EC_50_ values being even lower when compared with BSF427 (BSF 427 EC_50_: 0.489 µg/ml, AAC DKO EC_50_: 0.155 µg/ml). These results indicate that the AAC DKO cells rely on the F_o_F_1_-ATP synthase activity and maintain their ΔΨm through the reversed activity of this oligomycin-sensitive complex even in the absence of AAC.

### AAC DKO is unable to import ATP into the mitochondrial matrix

To examine whether there is an alternative way for the cytosolic ATP to cross the mitochondrial inner membrane in the absence of AAC, we assayed the capacity of the BSF427 and AAC DKO mitochondrion to build up ΔΨm by the proton-pumping activity of F_o_F_1_-ATP synthase in the presence of external ATP. We permeabilized the plasma membrane of the parasite with 4 µM digitonin and measured changes in Safranin O fluorescence upon addition of 1 mM ATP. Safranine O is a fluorescent dye that upon membrane potential-dependent uptake into the mitochondrion undergoes spectral change, which is monitored by a fluorimeter. The detected changes in fluorescence values are used to estimate ΔΨm [40]. The control BSF427 cells were able to build up and retain ΔΨm as evidenced by a decrease in safranine O fluorescence, which is completely reversed by addition of carboxyatractyloside (CATR), the inhibitor of AAC. Subsequent addition of oligomycin before the uncoupler SF 6847 had no further effect on depolarization (Figure 3A, black line). No changes in fluorescence were detected when the addition of CATR preceded that of ATP confirming that decrease in safranine O fluorescence is due to the activity of AAC (Figure 3A, red line). The AAC DKO were unable to generate ΔΨm in the presence of external ATP clearly showing that no ATP is able to enter the mitochondrial matrix. Importantly, a v5-tagged addback of AAC, expressed from a tubulin gene locus upon the addition of tetracycline, fully rescued the ability to polarize the inner membrane (Figure 3B). Therefore, the failure of the AAC DKO mutant to establish ΔΨm in permeabilized cells in the presence of exogenous ATP can be attributed specifically to the loss of AAC.

**Figure 3.**
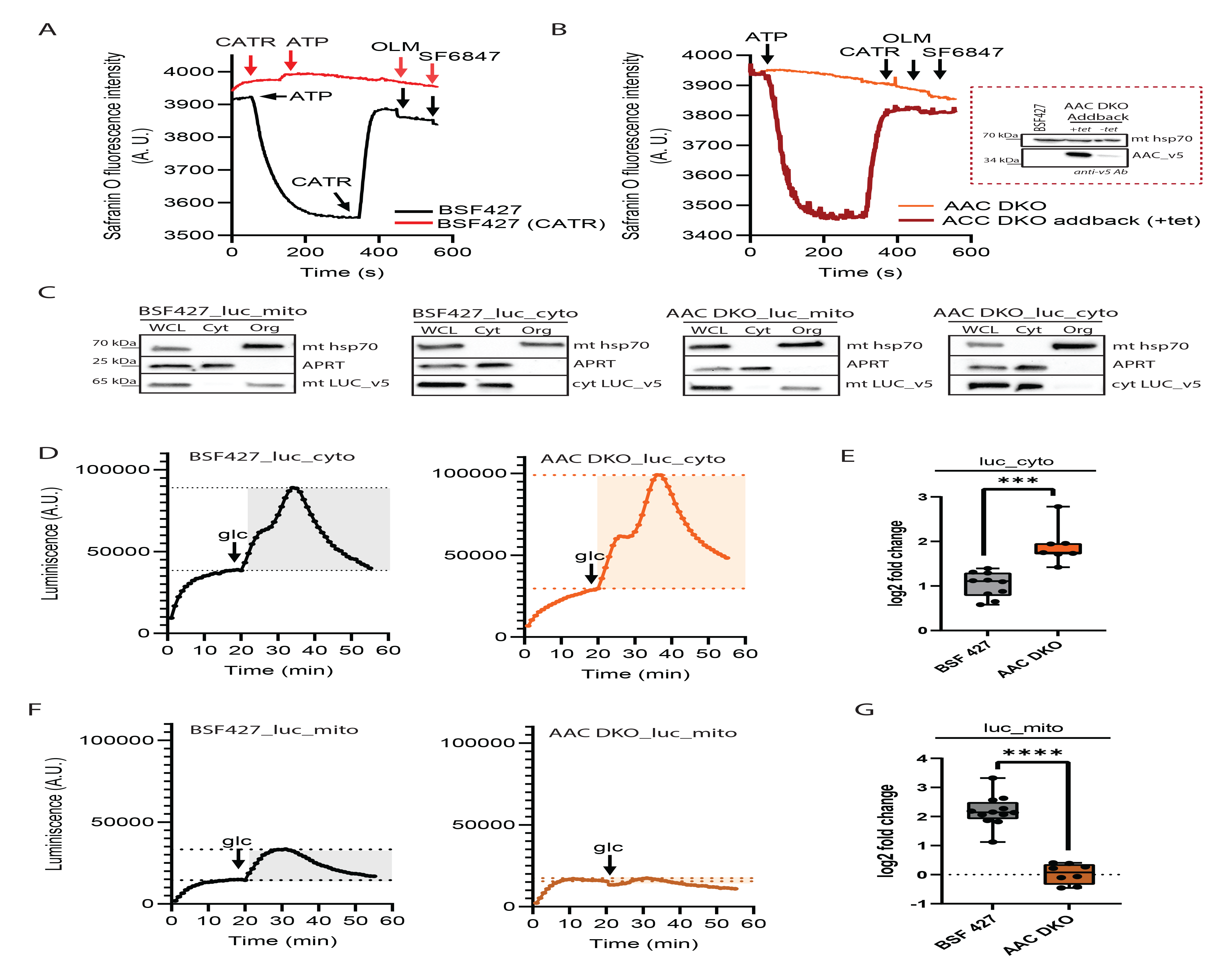
In the absence of AAC, the cells are unable to import cytosolic ATP to the mitochondrial matrix. A. Mitochondrial membrane polarization detected using Safranine O dye in digitonin-permeabilized BSF 427 cells in the presence of ATP. Carboxyatractyloside (CATR), the AAC inhibitor was added before the ATP (red line) as a control for no membrane polarization due to the inability to import ATP into the mitochondrial matrix. Oligomycin (OLM) was added after the CATR to induce depolarization. SF6847, an uncoupler, was added to test any further depolarization. ATP, CATR, OLM and SF 6847 were added where indicated. B. Mitochondrial membrane polarization detected using Safranine O dye in digitonin-permeabilized AAC DKO and AAC DKO Addback cells in the presence of ATPCATR, OLM and SF 6847 were added where indicated. The inset shows western blot analysis of BSF427, AAC DKO Addback cells grown in the presence or absence of tetracyline, probed with anti-v5 monoclonal antibody and anti-mt Hsp70 antibody as a loading control. C. Subcellular localization of v5-tagged luciferase without (luc_cyto) or with mitochondrial localization signal (luc_mito) endogenously expressed in BSF 427 and AAC DKO cells was determined in whole cell lysates and in the corresponding cytosolic and organellar fractions separated by digitonin extraction. Purified fractions were analyzed by Western blotting with the following antibodies: anti-v5, anti-mt Hsp70 (mitochondrial marker), and anti-adenosine phosphoribosyltransferase (APRT) (cytosolic marker). The relevant sizes of the protein marker are indicated on the left. D. Representative data of basal (first peak) and glucose-induced (second peak) levels of bioluminescence detected by a plate reader in the cytosol of BSF 427_luc_cyto (left pane) and AAC DKO_luc_cyto (right panel) using 25 µM luciferin. E. Quantification of changes in ATP levels upon 5mM glucose addition in BSF 427_Luc_cyto and AAC DKO_luc_cyto. Box and whiskers plots, n=7-10, *** p< 0.001 F. Representative data of basal (first peak) and glucose-induced (second peak) levels bioluminescence detected by a plate reader in the mitochondrial matrix of BSF 427_luc_mito (left pane) and AAC DKO_luc_mito (right panel) using 25 µM luciferin. G. Quantification of changes in ATP levels upon 5mM glucose addition in BSF 427_Luc_mito and AAC DKO_luc_mito. Box and whiskers plots, n=8-11, *** p< 0.001

To confirm that in the absence of AAC the cytosolic ATP is unable to enter mitochondrial matrix also in live cells, we generated reporter BSF427 and AAC DKO cell lines constitutively expressing mitochondrial or cytosolic firefly luciferase fused with C-terminal v5 tag. To ensure mitochondrial localization of the luciferase, its gene was fused with known mitochondrial localization signal of iron-sulphur cluster assembly protein ISCU [41]. The expression of the tagged luciferases and their appropriate localization in the cytosol or mitochondria were verified by western blotting (Figure 3C).

First, we monitored amounts of ATP in the cytosol in BSF427_luc_cyto and AAC DKO_luc_cyto cell lines. The expressed luciferase catalyzes oxidation of membrane-permeable D-luciferin producing a flash of light that is proportional to the amount of ATP present in the sample. Addition of glucose into the sample caused an increase in the cytosolic ATP levels in both cell lines suggesting an immediate glucose oxidation and generation of ATP by glycolysis (Figure 3D). Interestingly, the AAC DKO cells showed higher increase in ATP levels when compared to BSF 427 (Figure 3E), possibly because the ATP was not being drained to the mitochondrial matrix due to the lack of the AAC. Indeed, in the case of cell lines expressing mitochondrially localized luciferase, addition of glucose caused a spike in mitochondrial ATP levels in BSF427_luc_mito cells, but not in AAC DKO_luc_mito (Figure 3F, 3G). This result suggests that no glucose-derived ATP was imported to the mitochondrial matrix in the absence of AAC.

### AAC DKO is more sensitive to methyltriphenylphosphonium (TPMP), an inhibitor of **α**- ketoglutarate dehydrogenase

The ability of AAC DKO mutant to maintain the ΔΨm despite its inability to import cytosolic ATP to the mitochondrial matrix may suggest an intramitochondrial source of ATP. Since BSF trypanosomes are devoid of oxidative phosphorylation, this ATP can be generated by substrate-level phosphorylation, with the ATP-producing enzyme succinyl-CoA synthetase (SCoAS) being the best candidate (Figure 1). To determine if any of the mitochondrial enzymatic pathways that leads to generation of succinyl-CoA, the substrate for SCoAS, are affected by the lack of AAC, we performed label-free quantitative proteomic analysis. The BSF427 and ACC DKO cell lysates were analyzed in quadruplicates, measured with a 4-hour liquid chromatography gradient on a high-resolution mass spectrometry and data were analyzed by MaxLFQ. We quantified 3654 protein groups with a minimum of 2 peptides (1 unique) and present in at least two out of four replicates. Overall, the expression of only 76 proteins was significantly downregulated in AAC DKO, most of which were hypothetical and ribosomal proteins. A total of 44 proteins were significantly upregulated (p < 0.05, Figure 4A, Supplemental Table S1). Focusing specifically at the mitochondrial enzymes involved in the oxidative metabolism of glucose-derived pyruvate, threonine, and glutamine/glutamate, we found that the expression of these enzymes was only slightly affected by the absence of AAC (e.g. malic enzyme (ME)), with the exception of succinate dehydrogenase subunit 1 (SDH1), isocitrate dehydrogenase (IDH), and alanine aminotransferase (AAT), which were upregulated more than ∼1.5 times. In addition, some subunits of PDH, α-ketoglutarate dehydrogenase, branched-chain ketoamino acid dehydrogenase and mitochondrial pyruvate carrier 2 (MPC2), were also upregulated, but they did not meet our statistical criteria (Figure 4A, Supplemental Table S1). This data suggests that the AAC DKO mutant does not undergo major restructuring of its global proteomic landscape in response to the absence of AAC, but certain mitochondrial metabolic activities may be upregulated. However, our metabolomic analysis revealed no significant changes in the relevant metabolites (Figure 4B, supplemental Table S2) except for a slight increase in 2-hydroxyethylthiamin diphosphate (2HE-ThPP, ca 1.5 times, p-value = 0.02), a specific intermediate in pyruvate conversion to acetyl-CoA by PDH that corroborates the upregulation of this pathway toward the mitochondrial ATP synthesis. We also detected a slight accumulation of ATP (1.4 times, *p-*value = 0.02) and GTP (1.7 times, *p-*value = 0.008) at the cellular level in the AAC DKO mutant. These results suggest that deletion of AAC genes is not associated with major changes in the steady-state metabolism of *T. brucei* bloodstream forms.

**Figure 4.**
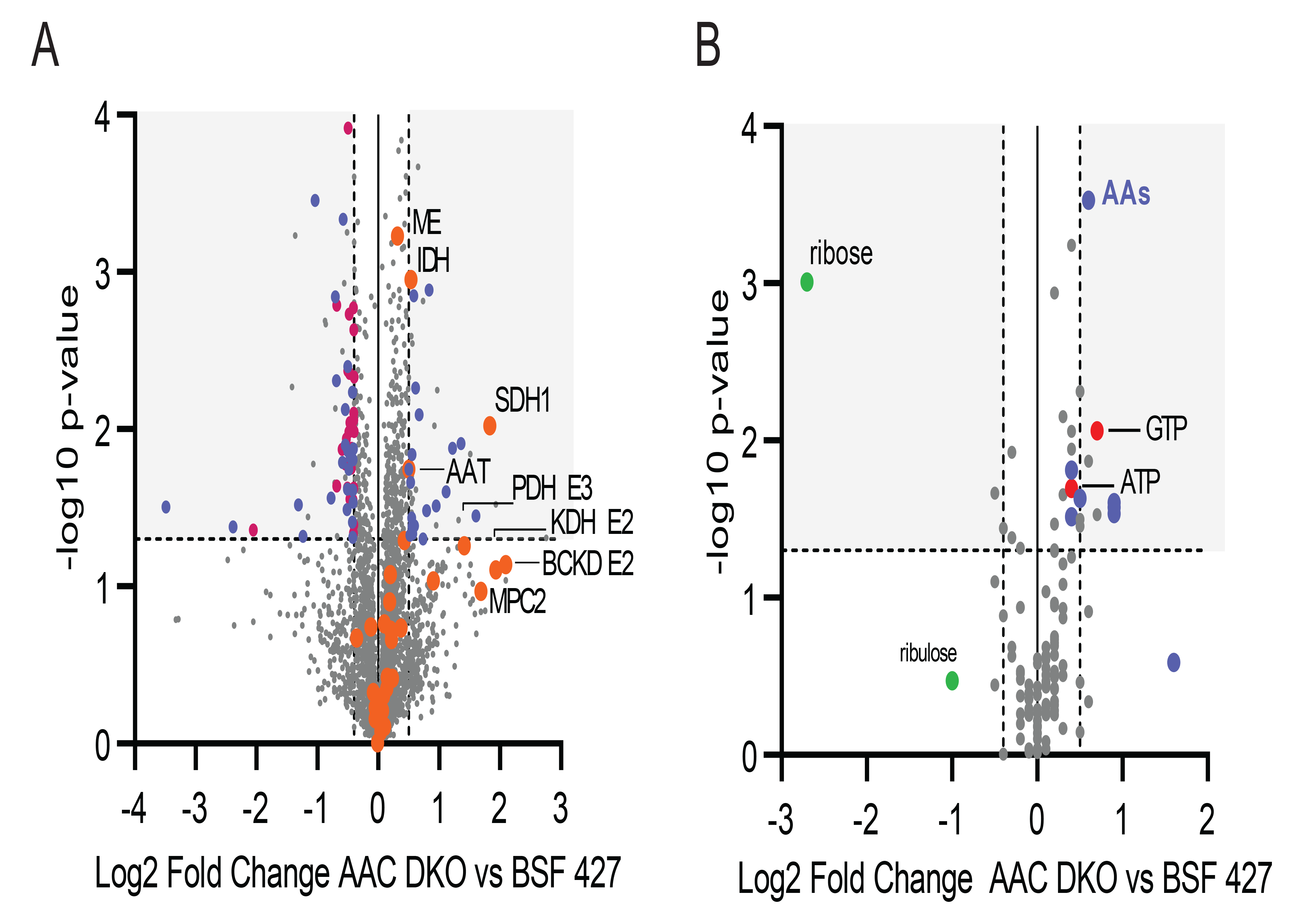
Proteomic and metabolomic profiling of AAC DKO cells. A. Volcano plots showing a comparison of protein expression levels (3654 protein groups) between BSF427 and AAC DKO cells. Log2 fold change values of averaged LFQ intensities from quadruplicate experiments are plotted against the respective −log10- transformed *P* values. Significantly changed hypothetical proteins are shown in blue, down-regulated cytosolic ribosomal proteins are shown in dark red. Mitochondrial enzymes involved in amino and keto acid oxidation including TCA cycle enzymes are highlighted in orange. ME, malic enzyme, IDH, isocitrate dehydrogenase, SDH1, succinate dehydrogenase subunit 1, AAT, alanine aminotransferase, PDH E3, subunit of pyruvate dehydrogenase, KDH E2, subunit of α-ketoglutarate dehydrogenase, BCKD E2, subunit of branch chain keto acid dehydrogenase, MPC2, mitochondrial pyruvate carrier 2, MCP14, mitochondrial carrier protein 14. B. Volcano plot showing the detected metabolites (124 metabolites) analyzed in BSF 427 and AAC DKO cells. Log2 fold change values of the average of mean peak area from quadruplicate experiments are plotted against the respective −log10 transformed *P* values. AAs, amino acids

Because changes in the levels of proteins or metabolites do not provide information about the importance of the mitochondrial metabolic pathways in question under specific conditions, we tested the importance of pyruvate, threonine, and α-ketoglutarate dehydrogenases for the cell viability using an Alamar Blue assay and their known inhibitors. We found no differences in the sensitivity of BSF427 and AAC DKO cells to sodium arsenite (BSF 427 EC_50_ = 0.22 µM vs AAC DKO EC_50_ =0.19 µM), an inhibitor of PDH, and quinazolinecarboxamide compound QC1 (BSF 427 EC_50_ = 11.3 µM vs AAC DKO EC_50_ = 9.7 µM) or tetraethyl thiuram disulphide (TETD) (BSF 427 EC_50_ = 9.9 µM vs AAC DKO EC_50_ = 9.4 µM), inhibitors of TDH [42].

The two separate metabolic pathways that include PDH and TDH and lead to generation of acetyl-CoA from pyruvate and threonine, respectively, were shown to be complementary to each other [31]. This could explain why no effect on sensitivity towards these inhibitors was detected. Importantly, the AAC DKO mutants were 18-fold more sensitive to the treatment with methyltriphenylphosphonium chloride (TPMP) (Figure 5A), a compound that has an inhibitory effect on α-ketoglutarate dehydrogenases [43]. TPMP-derived EC_50_ returned to the control levels of BSF427 in cells expressing a v5-tagged AAC addback copy (Figure 5A) confirming the specificity of the observed phenotype to the lack of AAC. Inhibition of α-ketoglutarate dehydrogenase leads to lowered production of succinyl-CoA, the substrate for the ATP-producing SCoAS. Interestingly, the phenotype was enhanced six-fold in cells in which ASCT expression was suppressed in the background of AAC DKO (Figure 5B and C). ASCT converts the pyruvate-or threonine-derived acetyl-CoA to acetate by coupling its transferase activity with ATP-producing SCoAS activity (Figure 1). The higher sensitivity of AAC DKO/ASCT RNAi cells to TPMP highlights the importance of the mitochondrial substrate phopshorylation for the AAC DKO cells. Our results suggest that in the absence of AAC, the null mutant is more dependent on the mitochondrial activities, which lead to generation of acetyl-and succinyl-CoA substrates feeding the two substrate phosphorylation pathways linked by the activity of SCoAS (Figure 1).

**Figure 5.**
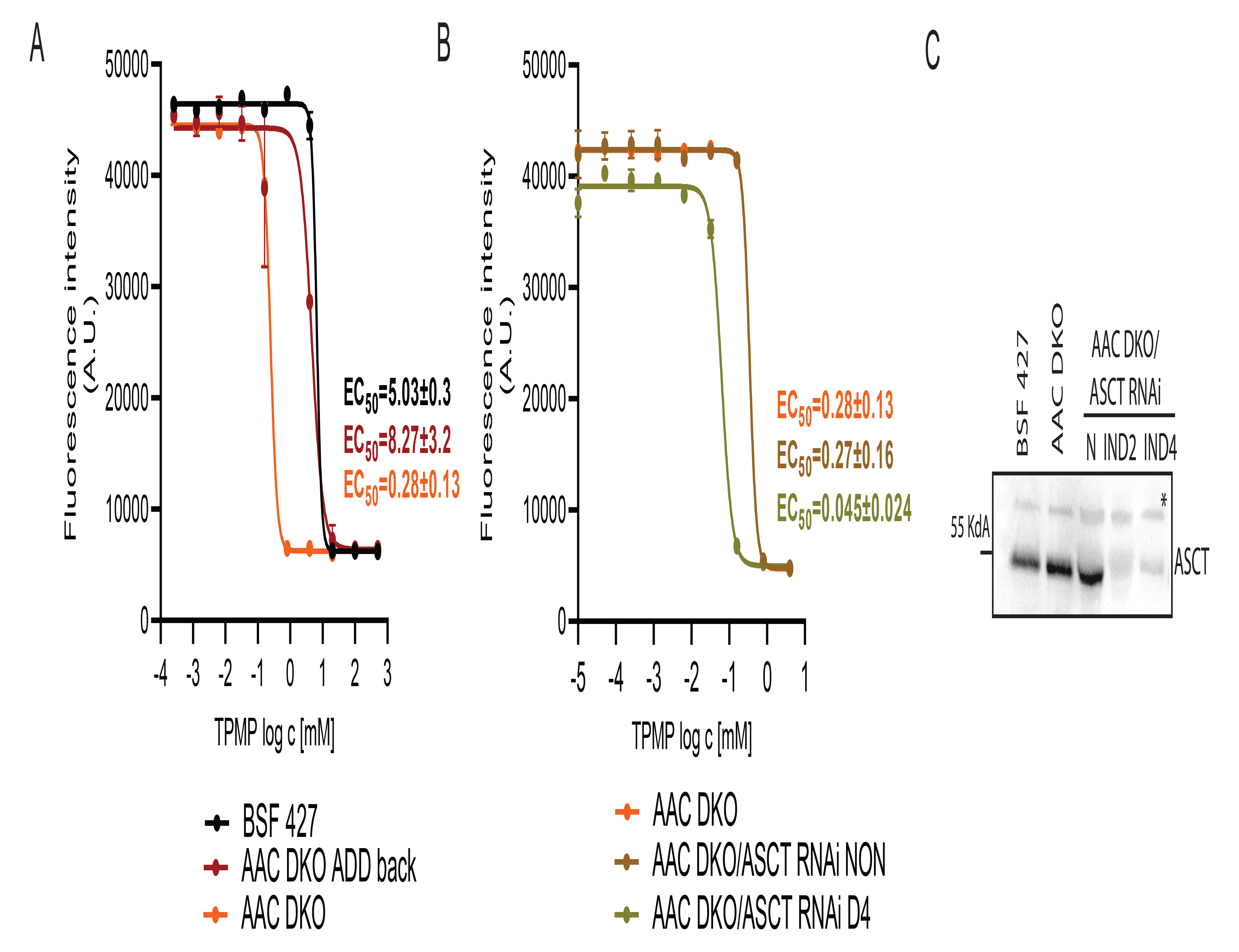
The AAC DKO cells are more sensitive to the treatment by TPMP, an inhibitor of α-ketoglutarate dehydrogenase. A. Sensitivity of BSF 427, AAC DKO, AAC DKO_addback to TPMP estimated by Alamar blue cell viability assay. B. AAC DKO/ASCT RNAi noninduced (NON) and cells induced for 4 days (D4) to TPMP estimated by resazurine cell-viability assay. The dose-response curves were calculated using GraphPad Prism 8.0 software. The calculated EC_50_ values are shown in graphs and are expressed in mM. C. Western blot analysis of BSF427, AAC DKO and AACDKO/ASCT RNAi cells uninduced and induced for 2 and 4 days using anti-ASCT antibody. *-non-specific band serving as a loading control.

### SCoAS is expressed and active in the BSF cells and its deletion leads to lowered virulency of the parasites in mice model

To assess the importance of SCoAS to the BSF parasites, we generated a double knockout of the β subunit of SCoAS (Tb927.10.7410) in HMI-11 medium (Figure 6A). Replacement of both SCoAS alleles with resistance markers was confirmed by PCR (Figure 6B), and the absence of the gene product was confirmed by Western blot using specific antibody raised against recombinant SCoAS (Figure 6C). The SCoAS enzyme was localized to the mitochondrial matrix as expected (Figure 6D). We also developed a specific enzymatic assay for SCoAS activity that encompasses the two substrates of this enzyme, succinyl-CoA and ADP, and produces ATP and CoA, the latter being tracked colorimetrically. The soluble mitochondrial fraction isolated from the control and AAC DKO cells showed comparable enzyme activity, whereas no SCoAS activity was detectable in the SCoAS DKO cells, confirming the specificity of this assay and the absence of alternative gene encoding the β subunit of SCoAS (Figure 6E). The SCoAS DKO mutants were viable in both HMI-11 and CMM media (Figure 6F and 6G). To investigate whether SCoAS is essential for the establishment of infection in animals, we inoculated groups of seven mice with BSF 427 and SCoAS DKO cells. Mice infected with the control parasites died 5-6 days after infection or had to be euthanized for ethical reasons when a parasitemia of 10^8^/ml was reached. In the case of SCoAS null mutants, four of the infected mice were not sick after two weeks and three survived the infection (Figure 6H). The SCoAS DKO addback cell line (western blot confirmed, Figure 6I) expressing SCoAS from the tubulin locus was again fully virulent and behaved the same as the control cells, confirming that the virulence defect was specifically due to loss of SCoAS (Figure 6J).

**Figure 6.**
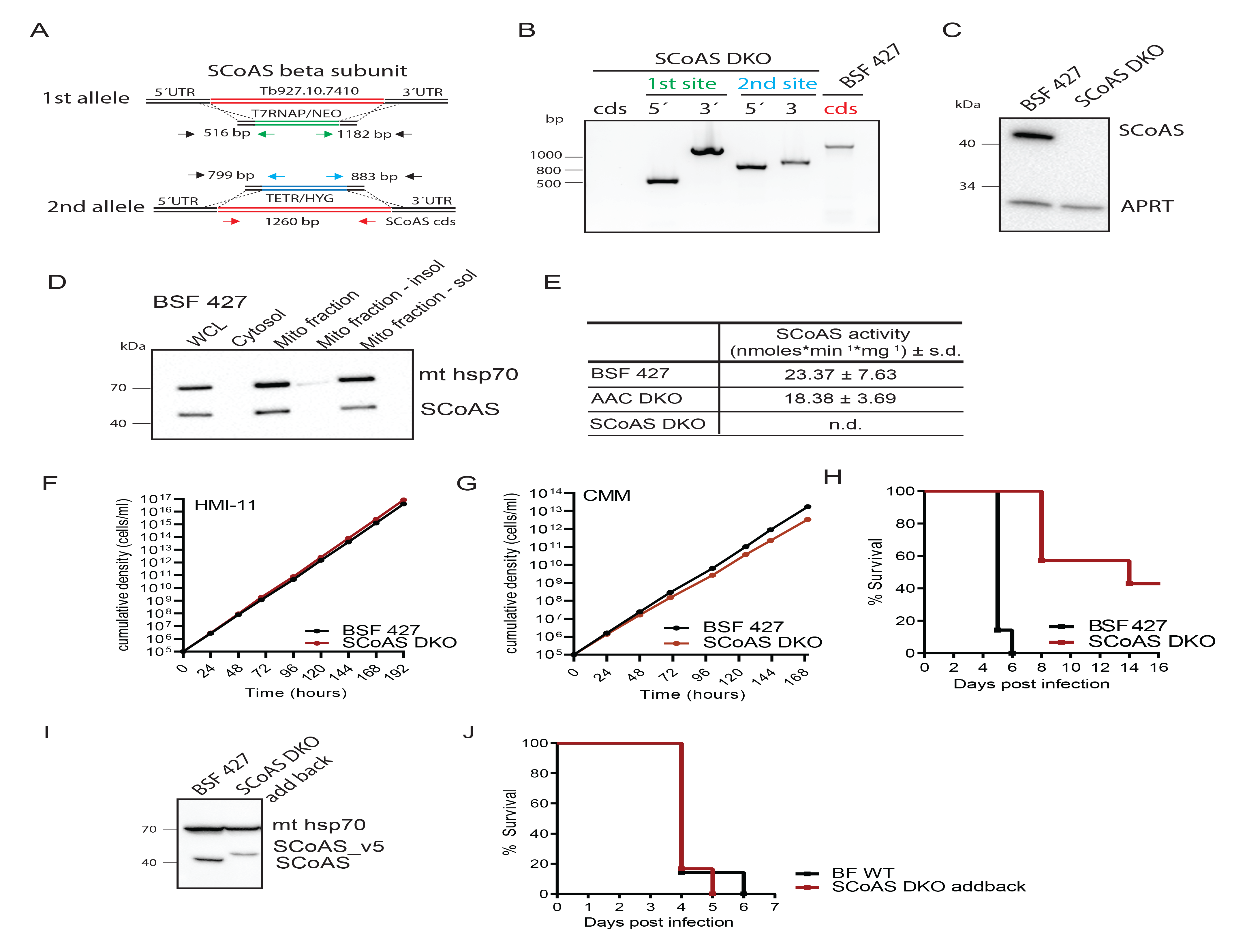
SCoAS DKO cells are viable in vitro but exert lower virulence in animal model. A. The strategy to generate SCoAS DKO involved replacement of both alleles with resistance genes conferring neomycin and hygromycin resistance. B. PCR verification for the elimination of both SCoAS alleles in SCoAS DKO cell line C. Immunoblot analysis of SCoAS DKO cells using specific anti-SCoAS antibody. Immunodetection of cytosolic APRT served as a loading control. D. Subcellular localization of SCoaS using BSF 427 cells. WCL, whole cell lysate, Cyt, cytosol, Mito, mitochondrial, insol, insoluble, sol, soluble E. Enzymatic activity of SCoAS measured in mitochondrial lysated extracted from BSF 427, AAC DKO and SCoAS DKO cells. F. Growth of AAC DKO cells compared to wild-type BSF 427 in HMI-11 and CMM medium measured for at least 7 days. G. The survival rate of 7 female BALB/c mice which were intraperitoneally infected with SCoAS DKO and wild-type BSF 427 parasites. The infected mice were monitored for 14 days. H. The survival rate of 7 female BALB/c mice which were intraperitoneally infected with SCoAS DKO Addback and wild-type BSF 427 parasites. The SCoAS DKO Addback infected mice were supplied with water containing doxycycline to induced expression of the addback SCoAS copy. The mice were monitored for 6 days. I. Immunoblot analysis of BSF 427 and SCoAS cDKO cell line inducibly expressing v5-tagged SCoAS using specific anti-SCoAS antibody. Immunodetection of mitochondrial hsp70 served as a loading control. J. The survival rate of 7 female BALB/c mice which were intraperitoneally infected with BSF 427 and SCoAS DKO_addback parasites.

### Metabolomic analysis of SCoAS mutants reveals changes in the levels of relevant metabolite

To identify possible metabolic changes in SCoAS DKO trypanosomes at the protein level, we performed quantitative label-free proteomic analyses of SCoAS DKO whole cell lysates and compared them with BSF 427 samples. Among the 3655 proteins identified by at least two peptides, only 17 and 21 proteins were up-and down-regulated by more than 1.5-fold, respectively, in the mutant cell line (p < 0.05), corresponding to approximately 0.5% of the proteome. The SCoAS subunit α was downregulated possibly in response to lack of its binding partner the SCoAS subunit β. Due to the small size of the significantly altered hits, the GO ontology enrichment analyzes did not reveal any enrichment of GO term categories (Figure 7A, Table S1).

**Figure 7.**
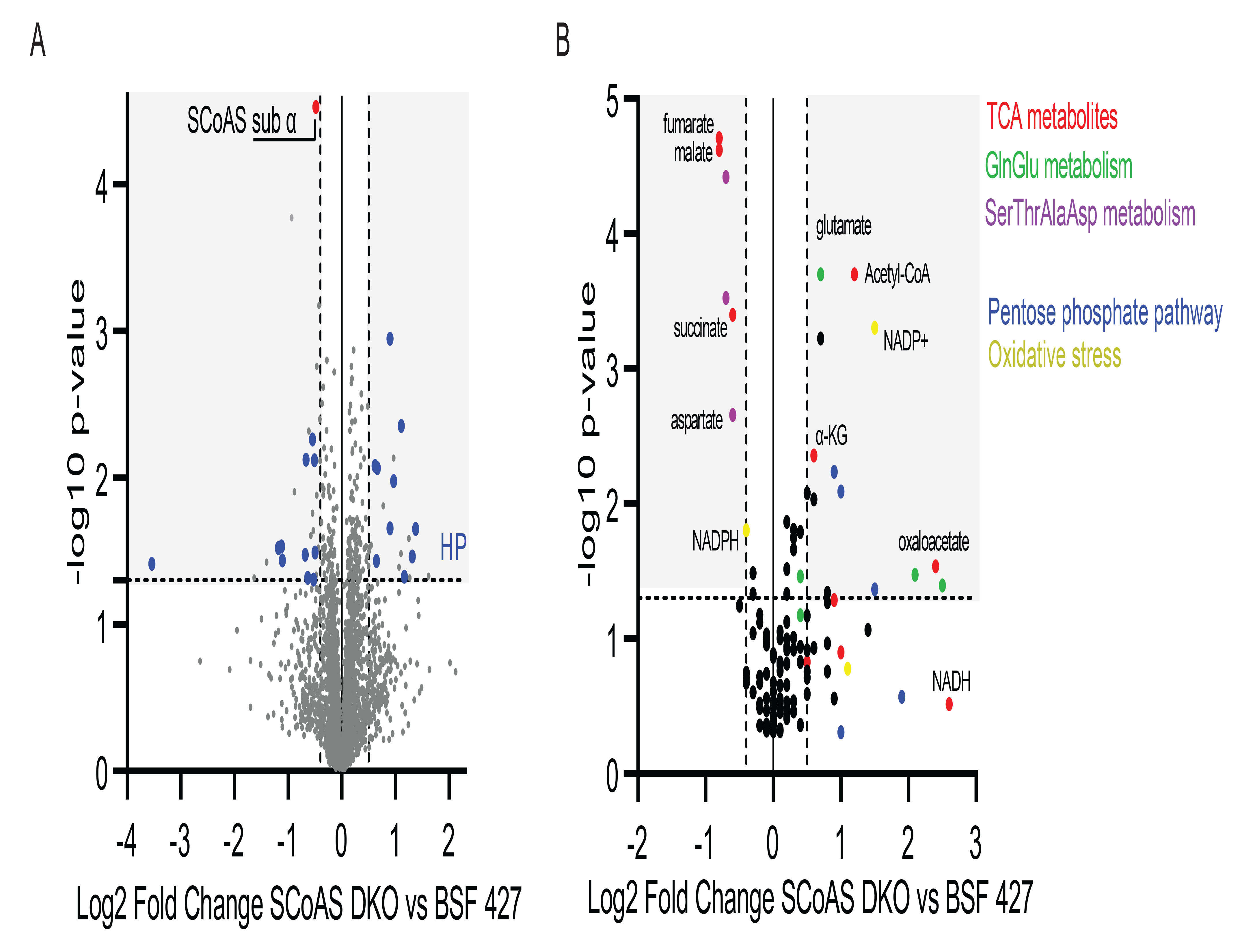
Proteomic and metabolomic profiling of SCoAS DKO cells. A. Volcano plots showing a comparison of protein expression levels (3654 protein groups) between BSF427 and SCoAS DKO cells. Log2 fold change values of averaged LFQ intensities from quadruplicate experiments are plotted against the respective −log10- transformed *P* values. Significantly changed hypothetical proteins are shown in blue. SCoAS sub α, subunit α of SCoAS α/β complex B. Volcano plot showing the detected metabolites (125 metabolites) analyzed in BSF 427 and AAC DKO cells. Log2 fold change values of the average of mean peak area from quadruplicate experiments are plotted against the respective −log10 transformed *P* values. Metabolites derived from the reaction of TCA cycle, glutamine/glutamate metabolism, serin/threonine/alanine/aspartate metabolism, pentose phosphate pathway and oxidative stress are highlighted in red, green, purple, blue and yellow respectively. α- KG, α-ketoglutarate.

We also performed targeted metabolomic analysis of the mutant and control parasites. Of the 127 metabolites identified, we found an enrichment of acetyl-CoA and α-ketoglutarate, the metabolites upstream of the substrate phosphorylation pathways, whereas succinate, malate, and fumarate, the downstream metabolites, were strongly downregulated. In addition, the increased levels of glutamate and oxaloacetate and the decreased levels of aspartate points to the activity of mitochondrial aspartate aminotransferase, an enzyme that is expressed in BSF cells. Alterations were also observed in metabolites belonging to the pentose phosphate pathway, the TCA cycle, and amino acid metabolism (Figure 7B, Supplementary Table S3).

To determine whether the absence of SCoAS also affects the excretion of metabolic end products, we compared BSF 427 and SCoAS DKO cells incubated in uniformly [U-^13^C]- enriched glucose-containing PBS by ^1^H NMR spectrometry. As expected, BSF cells excreted predominantly high amounts of pyruvate derived from glucose (79.8%), alanine (10.6%), lactate (4.1%), acetate (3.9%) and succinate (1.6%) (Figure 8A, left panel). Analysis of SCoAS DKO mutant revealed that there were no significant changes in glucose-derived excretion of pyruvate (82.8%), alanine (10.7%), succinate (1.2%), and lactate (5.0%). However, acetate excretion from glucose was completely abolished (Figure 8B, left panel). Because acetate can also be formed from threonine and ^1^H NMR spectrometry is able to distinguish ^13^C-enriched molecules from ^12^C molecules, we also performed a metabolite profiling assay with equal amounts (4 mM) of uniformly [^13^C]-enriched glucose and unenriched threonine to distinguish the metabolic origin of the end products (Figure 8A and 8B, middle and right panel). Interestingly, acetate excretion from both [U-^13^C]-enriched glucose and unenriched threonine was almost abolished, with only residual amounts of threonine-derived acetate, confirming that the ASCT/SCoAS cycle coupled to ATP generation is the primary source of acetate that is mainly excreted. Because SCoAS DKO cells did not exhibit a growth phenotype in either HMI-11 or CMM medium, we suggest that the activity of ACH in SCoAS DKO cells maintains intracellular acetate levels necessary for de novo biosynthesis of fatty acids in the absence of SCoAS [3], which is consistent with the residual amounts of acetate produced from threonine in the SCoAS DKO cells (Figure 8B). We also analyzed AAC DKO parasites for changes in glucose-, and threonine-derived metabolic fluxes, but no differences were found compared with the parental cells (Figure 8C, Supplementary Table S4).

**Figure 8.**
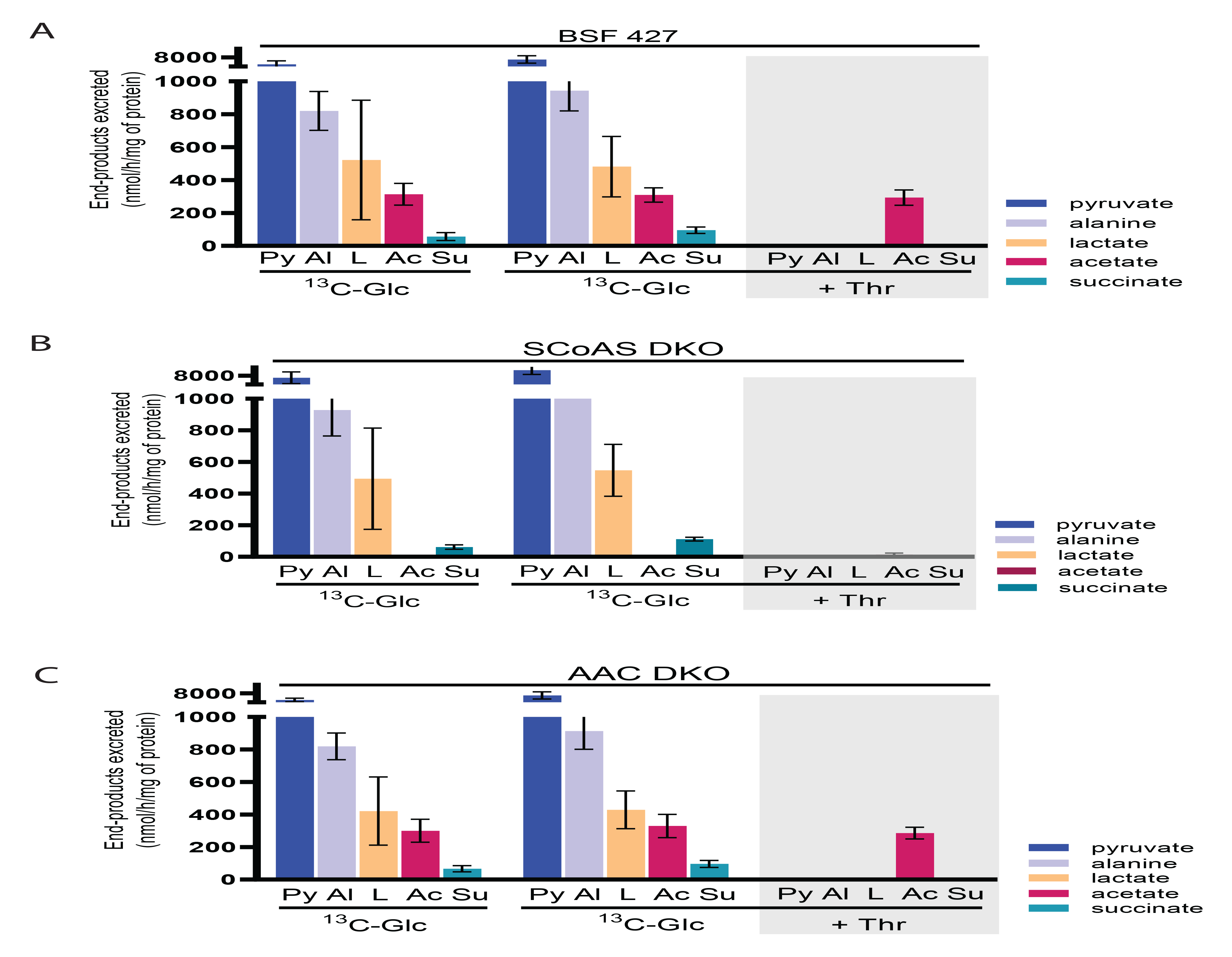
SCoAS DKO parasites do not excretes acetate. A, B, C. Proton (^1^H) NMR analyses of end-products excreted from the metabolism of ^13^C-enriched glucose. BSF 427 (A), SCoAS DKO (B) and AAC DKO (C) trypanosomes were incubated for 2.5 h in PBS containing 4 mM [U-13C]-glucose in combination with threonine (+Thr) or α-ketoglutarate (+α-KG) before analysis of the spent medium by ^1^H-NMR spectrometry. The amounts of each end-product excreted are documented in Table S3. Abbreviations: Ac, acetate; Al, alanine; L, lactate; Py, pyruvate; S, succinate.

Considering the significant changes in endo-and exo-metabolite profiles in SCoAS DKO cells, we set out to determine mitochondrial ATP levels in SCoAS DKO and compare them with BSF427 and AAC DKO cells. For this purpose, we also generated SCoAS DKO cells expressing v5-tagged luciferase with mitochondrial localization signal. The expression and localization of the mitochondrial luciferase was verified as for the mitochondrial luciferase-expressing BSF 427 and AAC DKO cells (Figure 2C and 9A). In all three cell lines, the expression of mitochondrial luciferases was at the comparable levels without statistically significant differences which allowed us to compare mitochondrial ATP levels between the different cell lines (Figure 9B and C). We monitored the luminescence emission that is proportional to the intracellular steady state ATP levels in the buffer without any carbon source. This is different from the experiments in Figure 2D - G where we measured the dynamics of glucose-induced ATP production. Here, the reaction was initiated only by addition of D-luciferin, the light emission rapidly increased and reached a plateau after certain time depending on the cell line (Figure 9D). The luminescence emissions at the plateau for each of the cell line from numerous independent experiments were plotted as a column graph (Figure 9E). The mitochondrial ATP levels in BSF427 and AAC DKO reached similar levels. Knowing that the mitochondrion of AAC DKO is not capable of importing ATP, this ATP pool must be produced intramitochondrially. Importantly, statistically less ATP was detected in the mitochondrial matrix of the SCoAS DKO mutant cells, when compared to AAC DKO and BSF427 cells. Interestingly, the detected lower mitochondrial steady-state ATP levels (Figure 9E) had no obvious effect on the maintenance of ΔΨm as no significant difference was detected in the fluorescence intensity of TMRE-stained BSF 427 and SCoAS DKO cell populations grown in either HMI-11 or CMM medium (Figure 9F). Furthermore, the ATP-induced polarization of the mitochondrial inner membrane in SCoAS DKO digitonin-permeabilized cells followed the same pattern as in BSF427 cells, suggesting that AAC is able to import ATP into the mitochondrion and this ATP is used to energize the membrane using F_o_F_1_-ATP synthase (Figure 9G and H).

**Figure 9.**
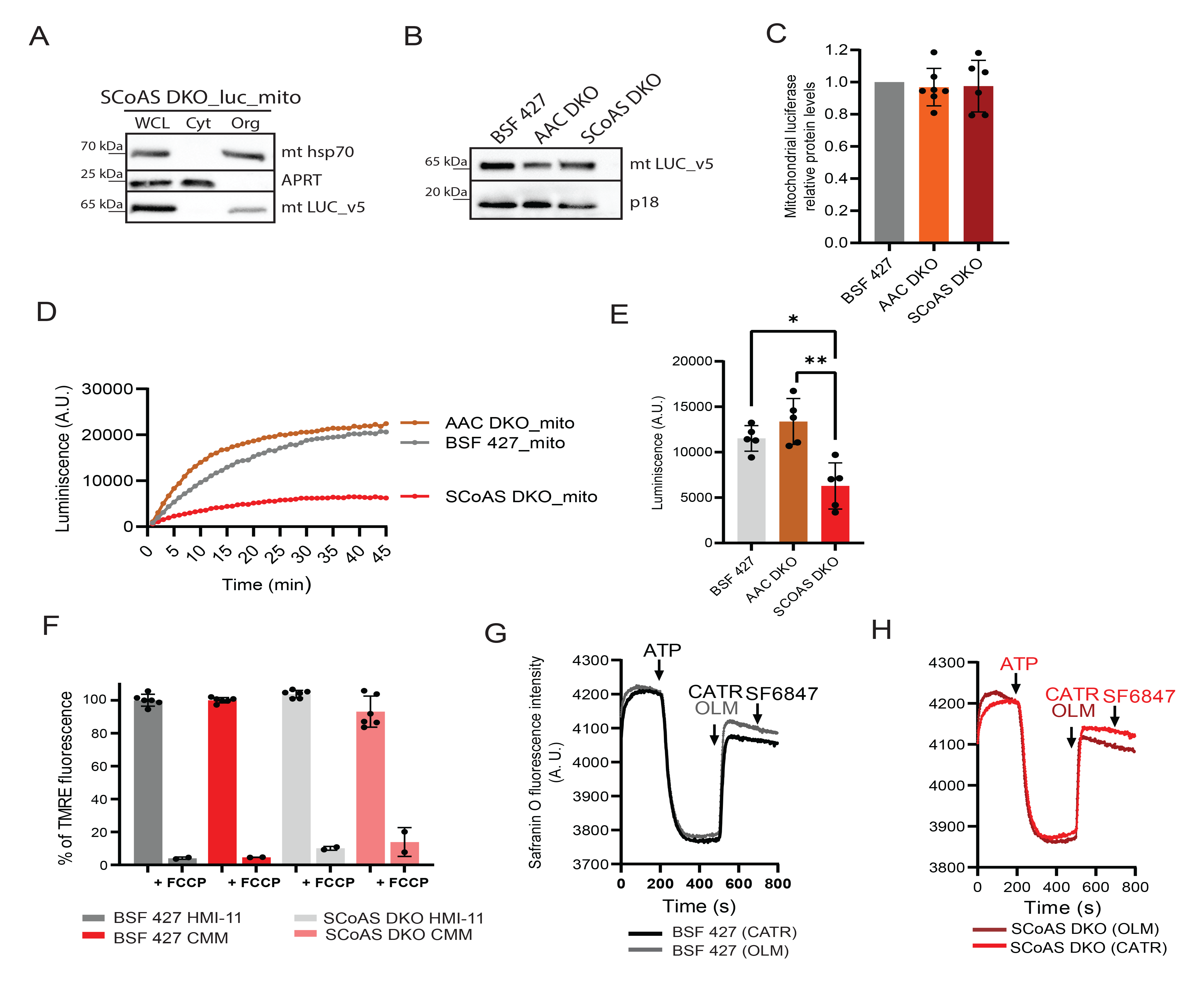
SCoAS DKO parasites have decreased mitochondrial ATP content, but are capable of ATP import and ATP hydrolysis. A. Subcellular localization of V5-tagged luciferase without (luc_cyto) or with mitochondrial localization signal (luc_mito) endogenously expressed in ScoAS DKO cells was determined in whole cell lysates and in the corresponding cytosolic and organellar fractions separated by digitonin extraction. Purified fractions were analyzed by Western blotting with the following antibodies: anti-v5, anti-mt Hsp70 (mitochondrial marker), and anti-adenosine phosphoribosyltransferase (APRT) (cytosolic marker). The relevant sizes of the protein marker are indicated on the left. B. Immunoblot of V5-tagged luciferase expressed in BSF 427_luc_cyto, BSF 427_luc_mito, AAC DKO_luc_cyto, AAC DKO_luc_mito, SCoAS DKO_luc_cyto, SCoAS DKO_luc_mito cells using antibodies against V5 tag. Antibody against subunit p18 of FoF1 ATP synthase was used as a loading control. C. The quantification analyses of luciferase expression in all cell lines by densitometry. The bars represent relative protein amounts of luciferase expression in AAC DKO and SCoAS DKO cells compared to luciferase expression in BSF 427. (mean, ± s.d., n= 6-7) D. Representative data of ATP measurements performed in living BSF 427_luc_cyto, BSF 427_luc_mito, AAC DKO_luc_cyto, AAC DKO_luc_mito, SCoAS DKO_luc_cyto, SCoAS DKO_luc_mito cells using 25 µM luciferin. E. Quantification of the luminescence measurement detected in BSF 427_luc_cyto, BSF 427_luc_mito, AAC DKO_luc_cyto, AAC DKO_luc_mito, SCoAS DKO_luc_cyto, SCoAS DKO_luc_mito. Graphs are derived from experiments which representative graphs are shown in D. (mean, ± s.d., n= 4-5, Student´s unpaired *t*-test, *< 0.05, **< 0.005). F. Flow cytometry analysis of TMRE-stained SCoAS DKO and BSF 427 cells grown in HMI-11 or CMM medium to measure ΔΨm. The addition of FCCP served as a control for ΔΨm depolarization (+FCCP). (means ± s.d., n= 6) G. Mitochondrial membrane polarization detected using Safranine O dye in digitonin-permeabilized BSF 427 cells (black/grey lines) and SCoAS DKO (light and dark red) in the presence of ATP. ATP, CATR, OLM and SF 6847 were added where indicated.

### SCoAS DKO cells are dependent on ATP import from the cytosol

In the absence of mitochondrial substrate phosphorylation pathways we tested if SCoAS DKO viability relies more on the AAC activity to import the necessary ATP into the mitochondrial matrix. The cell viability assay showed that SCoAS DKO cells are more sensitive to CATR and bongkrekic acid, known specific inhibitors of the AAC. The EC_50_ values of SCoAS DKO parasites were ∼25-fold lower in the case of carboxyatractyloside and ∼5-fold lower for bongkrekic acid when compared to BSF 427 (Figure 10A and 10B). Consistently with this observation, the ASCT DKO, which is defective in one of the two mitochondrial substrate phosphorylation pathways, showed only 10-fold higher sensitivity to CATR when compared to BSF 427 (Figure 10A). Tetracycline-induced expression of SCoAS in the background of the SCoAS null mutant restored the original EC_50_ values for CATR, confirming that the observed phenotype in CATR-sensitivity was due to the absence of SCoAS (Figure 10 C, D).

**Figure 10.**
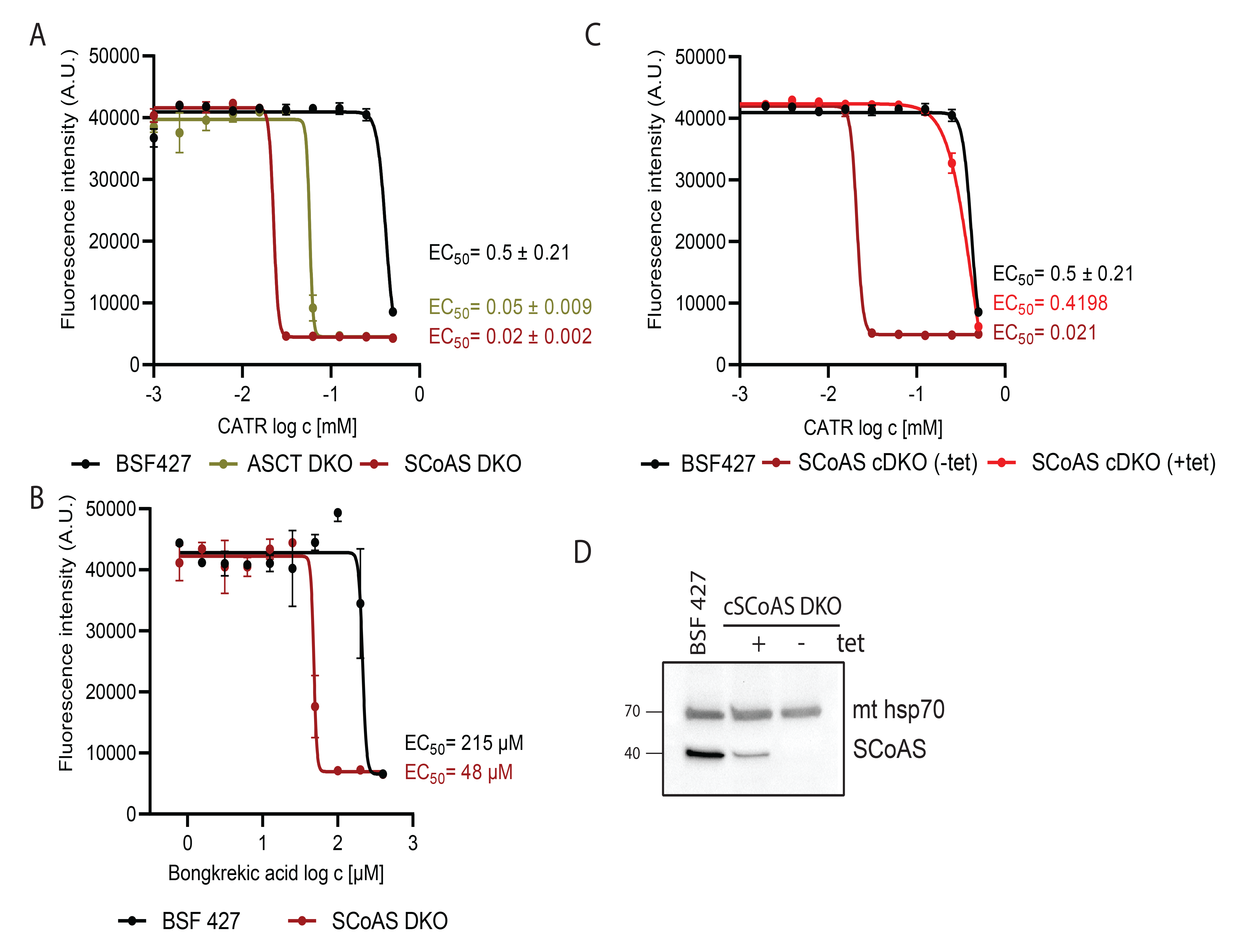
SCoAS DKO cells are more sensitive to CATR, an inhibitor of AAC. A. Sensitivity of BSF 427, SCoAS DKO, ASCT DKO to carboxyatractyloside (CATR) estimated by resazurine cell-viability assay. The dose-response curves were calculated using GraphPad Prism 8.0 software. The calculated EC_50_ values are shown in graphs and are expressed in mM. B. Sensitivity of BSF 427 and SCoAS DKO, ASCT DKO to bongkrekic acid estimated as in (A). C. Sensitivity of BSF 427, SCoAS cDKO noninduced (-tet) and 4-days induced (+tet) cells to carboxyatractyloside (CATR) estimated as in (A). D. Immunoblot of SCoAS cDKO noninduced (-tet) and 2-days induced (+tet) cells using SCoAS antibody. Immunodetection of mitochondrial hsp 70 served as a loading control.

In summary, it appears that *T. brucei* BSF parasites have two alternative options for mitochondrial ATP provision, intramitochondrial ATP production by substrate-level phosphorylation and ATP import from the cytosol *via* AAC. To test the essentiality of this intriguing functional interplay, we attempted to silence AAC expression by RNAi in the SCoAS DKO background and, conversely, to silence SCoAS expression in the AAC DKO background. Unfortunately, our numerous attempts failed. We were unable to select any prospering cell clones responding to tetracycline-induced silencing of the targeted gene. We explain this phenomenon by the background expression of dsRNA in the absence of tetracycline during the selection process causing a lethal phenotype. It is likely that these two pathways act complementarily to each other, and the absence of both pathways is not consistent with the survival of *T. brucei* parasites under the conditions used.

### Mitochondrial production of ATP by substrate-level phosphorylation is essential under glycerol-rich growth conditions

BSF trypanosomes can also utilize glycerol as the main carbon source [44, 45] although the growth rate is significantly reduced in glycerol-rich culture medium (CMM_gly) (Figure 11A). This is due to the limited capacity to metabolize glycerol compared to glucose, resulting in a slightly lower yield of cytosolic ATP when compared with cells grown in CMM_glc. Interestingly, BSF grown in CMM_gly excrete more acetate and succinate than those grown in CMM_glc suggesting higher activity of mitochondrial metabolic pathway [44]. Therefore, we further investigate the importance of SCoAS for cells grown in CMM_gly. The BSF 427 and AAC DKO cells were able to adapt to glycerol conditions well, albeit they grow at a slower growth rate. In contrast, SCoAS DKO was never able to establish an adapted dividing culture (Figure 11A). To investigate whether the reason for this observation is due to the essential role of mitochondrial substrate-level phosphorylations to maintain the ΔΨm under the conditions of lower cytosolic ATP yield in CMM_gly medium, we generated RNAi cells to silence SCoAS expression. In addition to HMI-11, SCoAS RNAi cells were adapted for growth in CMM_glc and CMM_gly. The efficiency of RNAi-mediated downregulation of SCoAS was verified under all three growth conditions by Western blot using specific antibodies (Figure 11B). The propagation of SCoAS RNAi was not affected by the addition of tetracycline to HMI-11 medium (Figure 11C). Furthermore, we did not detect any decrease in ΔΨm by flow cytometry in TMRE-stained noninduced and tetracycline-induced cells (Figure 11D). However, silencing of SCoAS in CMM_glc medium resulted in slower growth of the RNAi-induced cell population. Compared with BSF 427 cells, ΔΨm was decreased by approximately 30% in cells induced for 5 days, which most likely contributed to the mild growth phenotype of these cells. Most importantly, SCoAS RNAi cells grown in CMM_gly exhibited a severe growth phenotype associated with a sharp decrease in ΔΨm at days 1, 2, and 3 after induction. In this case, ΔΨm values fell below the minimum threshold required for *T. brucei* viability *in vitro* [21, 39]. Our data clearly show that the functional interplay between the AAC and an ATP-producing SCoAS activity depends on the environment. When cells encounter an environment with lower glucose concentration or different carbon source (i.e. glycerol), which yields lower cytosolic ATP levels or if they faced conditions that lead to higher consumption of the cytosolic ATP (e.g. stress), the BSF *T. brucei* parasites cannot compensate their mitochondrial substrate-level phosphorylation pathways by taping the cytosolic ATP pool and sustain the essential ΔΨm. The ability of AAC to reverse its activity depends on the levels on ΔΨm, cytosolic ATP levels and ATP/ADP ratio in mitochondrial matrix. Therefore, the parasite bioenergetics regulates the major contributing pathways of ATP provision that are fully compensatory when the parasite is in glucose-rich HMI-11 culture conditions. However, when *in vivo* or CMM_gly culture conditions our data show that the mitochondrial substrate phosphorylation pathways become more important for the parasite survival. Finally, we can conclude that BSF mitochondrion is capable of producing ATP.

**Figure 11.**
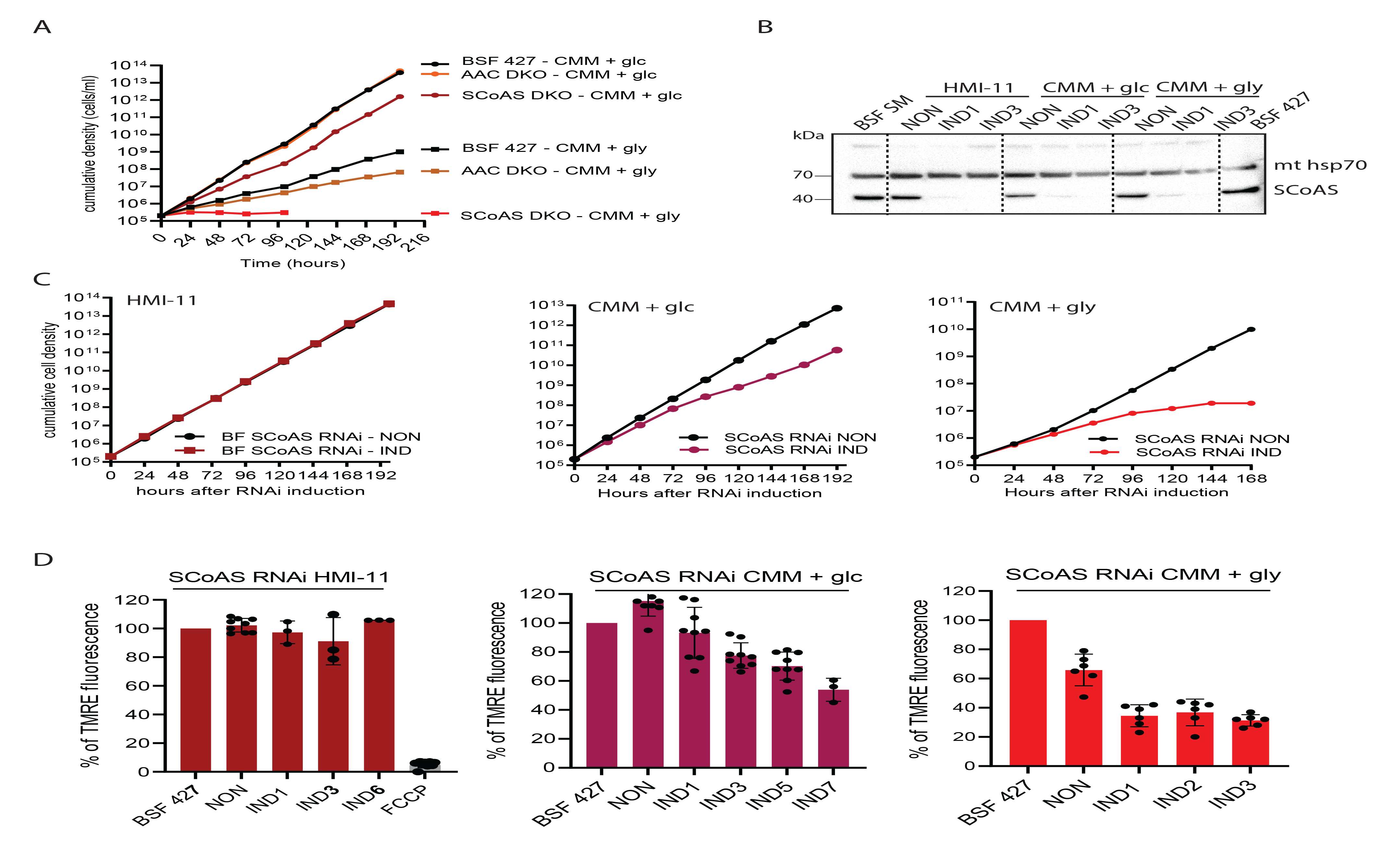
SCoAS RNAi silencing results in growth phenotype and decreased ΔΨm in CMM_glc and CMM_gly medium. A. Growth of BSF 427 and SCoAS DKO cells in HMI-11, CMM_glc and CMM_gly medium. B. Western blot analysis of whole cell lysates of SCoAS RNAi noninduced and induced (+tet) cells grown in HMI-11, CMM_glc and CMM_gly using antibodies against the SCoAS protein. The immunoblot probed with anti-mitochondrial hsp70 antibody served as loading controls. Glc, glucose, gly, glycerol. C. Growth of SCoAS RNAi noninduced (non) and tetracycline induced (IND) cells measured for 8 days in HMI- 11 (left), CMM_glc (middle) and CMM_gly (right). Glc, glucose, gly, glycerol. D. Flow cytometry analysis of TMRE-stained SCoAS RNAi noninduced and induced cells grown in HMI- 11 (right), CMM_glc (middle) and CMM_gly (left). (means ± s.d., n= 3- 9)

## Discussion

The widespread assumption of low mitochondrial activity of the *T. brucei* parasite has recently been challenged by proteomic and metabolomic data suggesting that certain metabolic pathways can be active if environmental conditions allow [46]. High flexibility and adaptability of the parasite organelle can be beneficial for the parasite when adapting to new host environments (e.g. when populating adipose tissues, skin) [47, 48]. Nevertheless, the historically established basic picture of the molecular mechanism maintaining ΔΨm is that proton-pumping F_o_F_1_-ATP synthase hydrolyzes ATP that is provided from the cytosol by the reverse activity of AAC. However, the mitochondrial acetate production pathway that is linked to an ATP-forming activity, is questioning this view [31, 32].

There is no doubt that F_o_F_1_-ATP synthase is the main molecular entity in the BSF *T. brucei* that is able to generate ΔΨm at the value between −150 to −180 mV [14, 15]. This electrochemical potential is then used for protein import and ion exchange. At the value of −150 mV, the cellular conditions (such as mitochondrial matrix ATP/ADP ratio, cytosolic ATP levels, etc.) allows the F_o_F_1_-ATP synthase to reach its reversal potential (E_rev_ATPase_). It was assumed that, under these conditions, most of the ATP feeding F_o_F_1_-ATP synthase is supplied by the AAC also operating in the reverse mode.

As described for BSF, mammalian cells can encounter conditions under which the F_o_F_1_-ATP synthase and AAC reach their reversal potentials [18]. Mathematical modeling of the mammalian cells capable of oxidative phosphorylation has shown that during mitochondrial membrane depolarization induced by ETC inhibition or hypoxia, the F_o_F_1_-ATP synthase reaches its E_rev_ATPase_ value first and opens a situation when AAC has not reversed yet, and ATP is therefore generated by mitochondrial substrate phosphorylation. If ΔΨm drops even further (i.e. shifts to less negative values), the AAC also reverses [18]. The AAC reversal potential (E_rev_AAC_) depends on many additional parameters, the most important ones being the ATP/ADP ratio in the mitochondrial matrix and ATP levels in the cytosol. In a situation where there is a lot of ATP in the mitochondrial matrix, the AAC has no reason to reverse. However, if the ATP/ADP ratio drops under certain value, the AAC starts importing ATP from the cytosol. The timing of this also depends on the cytosolic ATP levels, the higher the ATP levels, the sooner AAC reverses.

As a surprise came a finding that AAC DKO *T. brucei* BSF cells were viable *in vitro* and fully virulent in mouse model, suggesting that import of cytosolic ATP to the mitochondrion is dispensable. Further, our data clearly show that AAC is the only carrier that can import ATP into the mitochondrial matrix since high extracellular ATP concentrations (1 mM) did not induce mitochondrial membrane polarization in AAC DKO. Further, the addition of glucose to the live cells in defined buffer stimulated an ATP generation in the cytosol followed by an import of ATP into the mitochondrial matrix in BSF 427 cells, while in AAC DKO, no increase in mitochondrial ATP level was detected. This suggests that AAC must be present in order to import ATP molecules into the mitochondrial matrix.

Therefore, in the case of AAC DKO the culture medium and host environment provided enough nutrients to support mitochondrial ATP production by substrate phosphorylation pathways which are powerful enough to provide enough ATP to maintain ΔΨm at the levels compatible with the full viability and virulence. Although the metabolomic changes in AAC DKO may have pointed to some extent to higher activity of mitochondrial metabolic pathways linked to ATP production, the determined levels of metabolic end-products (i.e. pyruvate, acetate, succinate and alanine) showed no significant changes, suggesting that the lack of AAC did not cause a striking metabolic remodeling in order to adapt to its absence. This suggests that in the absence of AAC, the BSF mitochondrion is able to become fully independent of the cytosolic tap of ATP. This also suggests that the AAC is in a neutral position where it probably does not import ATP into the mitochondrial matrix at high efficiency (no striking phenotype in AAC DKO cells), but possibly it can reverse and thus initiate an efficient ATP import, whenever the mitochondrial substrate phosphorylation pathways are not able to provide enough ATP (Figure 12A). High levels of the cytosolic ATP allow for an immediate reversal of AAC and taping the cytosolic ATP pool to maintain ΔΨm. This is exemplified in SCoAS DKO, in which due to the lack of the ATP-producing activity of this enzyme, the mitochondrial ATP/ADP ratio is decreased. Indeed, our luciferase-based assay showed lower mitochondrial steady-state ATP levels when compared to BSF 427 and AAC DKO. Therefore, under the same value of ΔΨm as detected by the TMRE measurements, the AAC fully reverses imports ATP and compensate for the SCoAS loss (Figure 12C). Moreover, this activity now becomes important for the viability of the parasite as SCOAS DKO cells exert higher sensitivity to CATR, the inhibitor of AAC.

**Figure 12.**
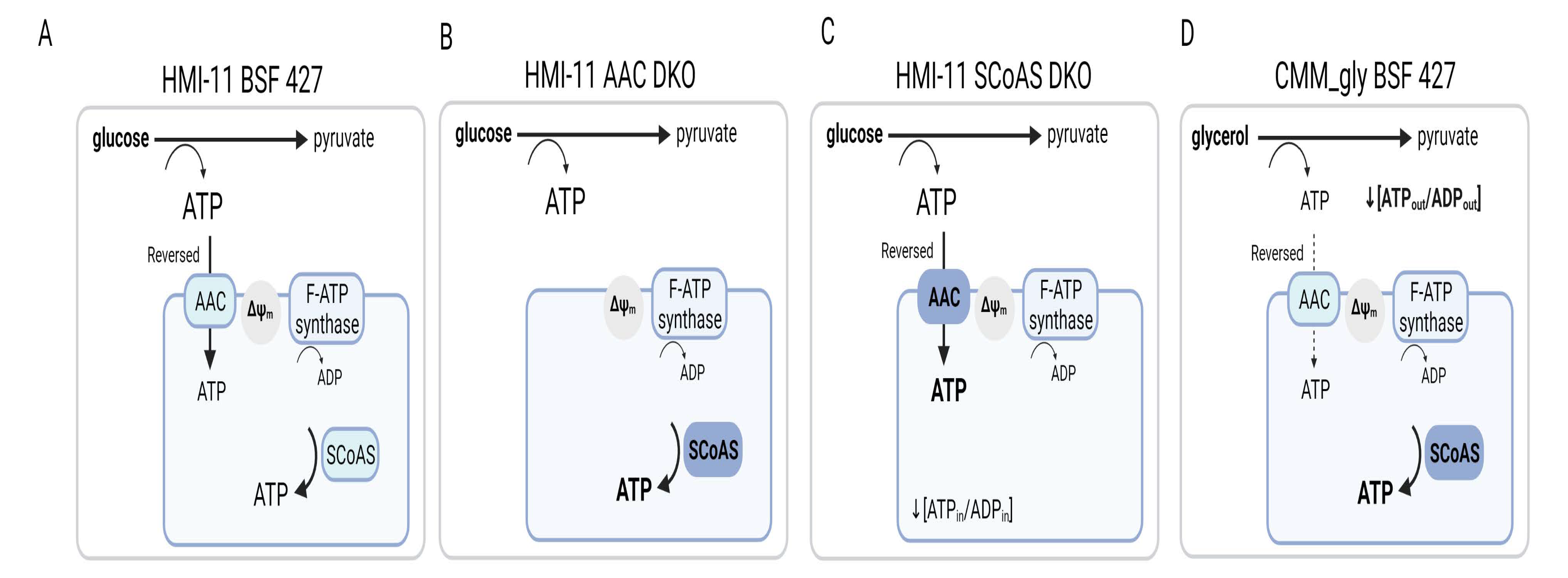
Schematic visualization of AAC and SCoAS activities interplay in BSF427 (A), AAC DKO (B) and SCoAS DKO (C) grown in HMI-11 and BSF 427 cultured in CMM_gly medium (D). AAC, ATP/ADP carrier, SCoAS, succinyl-CoA synthetase

Now the question arises where the substrate for the ATP-producing SCoAS, the succinyl-CoA, comes from. Based on the metabolic pathways mapped to the parasite mitochondrion [29-31], there are several options. First, succinyl-CoA is produced by ASCT enzyme from the pyruvate-and threonine-derived acetyl-CoA. The importance of the ASCT/SCoAS cycle for the ATP/ADP ratio in mitochondrial matrix is supported by the observation that ASCT DKO is 10- time more sensitive to the AAC inhibitor, CATR, compared to BSF 427. Originally, the metabolic pathways leading to the production of acetyl-CoA were studied from the point of acetate production, the essential precursor for de novo fatty acid biosynthesis [49]. Indeed, when both pathways leading to acetyl-CoA were genetically impaired, growth of BSF cells in HMI-11 medium is strongly affected because of the inability to produce acetate [31]. Interestingly, excretion of acetate was greatly reduced but not fully abolished in SCoAS DKO, suggesting that this baseline level of acetate production is sufficient to support fatty acid biosynthesis without effecting parasite growth rate. In agreement with this hypothesis, we estimated that ∼2% of acetate produced by insect form trypanosomes from glucose and threonine metabolism is incorporated into fatty acids, since among the ∼500 nmol of acetate excreted per h and per 10^8^ cells [4] only 10 nmol are incorporated into fatty acids per h and per 10^8^ cells [50]. This residual production of acetate in the SCoAS DKO is therefore due to the ACH activity [6].

Second potential source of succinyl-CoA is from α-ketoglutarate by the activity of α- ketoglutarate dehydrogenase, that is expressed in the BSF cells although its function for the parasite is so far enigmatic [51]. The α-ketoglutarate can be derived from glutamine, an amino acid that is consumed in significant amounts [29]. Another possible source of α-ketoglutarate are transamination reactions employing alanine and aspartate aminotransferases. There is no doubt about the activity of alanine aminotransferase, as BSF parasites excrete significant amounts of alanine from pyruvate. Interestingly, the alanine aminotransferase is probably essential for BSF cells [52], and thought to be a cytosolic enzyme, its localization was just recently challenged by Tryptag data that has placed this enzyme also into the mitochondrial matrix [53]. In addition, α- ketoglutarate should also be considered as an excellent potential external source of carbon, as recently observed for PCF trypanosomes [4].

Our metabolomic data of SCoAS DKO cells, which show a significant reduction of the SCoAS product succinate as well as the downstream metabolites fumarate and malate and the same time accumulation of the upstream metabolites acetyl-CoA, α-ketoglutarate and oxaloacetate, indicate a disruption of the mitochondrial metabolic network linked to SCoAS activity. However, it should be noted that our analysis does not differentiate between the glycosomal, cytosolic and mitochondrial pools of dicarboxylic acids, although it seems unlikely that the SCoAS absence would somehow affect the glycosomal reduction pathway to succinate, which functions mainly as a redox balancer for glycolysis. Nevertheless, further metabolomic studies using labeled-carbon sources (e.g. glucose, glutamine) are needed to determine the precise origin and localization of these metabolites.

Surprisingly, the growth of the SCoAS RNAi cell line in CMM medium was significantly affected while that of the SCoAS DKO cells was not. The growth defect of the RNAi SCoAS induced cells is probably due to a slightly lower ΔΨm, that is not observed for the the SCoAS DKO cells. It should be noted that the SCoAS DKO was subjected a step wise adaptation, by first deleting one allele and then growing the null mutant during 14-day before being analyzed, which is probably sufficient for the mutant to adapt its metabolism to the absence of SCoAS, while, the tetracycline-induced SCoAS RNAi cell line had to cope with a suddenly decreased expression of SCoAS.

In conditions when glycerol is the main carbon source, the BSF parasites metabolized it to pyruvate, alanine, acetate and succinate, as observed for glucose. To produce similar amounts of cytoplasmic ATP, twice as much glycerol (three-carbon compound) must be metabolized as glucose (six-carbon compound). The 427 BSF strain consumes only 1.5-times more glycerol than glucose when grown in CMM_gly and CMM_glc, respectively, explaining the significant growth delay observed in CMM_gly [44]. Interestingly, the absolute amounts of acetate produced in CMM_gly and CMM_glc is similar (283 *versus* 262 nmol/h/10^8^ cells [44], suggesting that maintaining mitochondrial substrate level phosphorylation is important, which is particularly relevant when cytosolic ATP is reduced, as expected in CMM_gly. This hypothesis is in agreement with the observation that the SCoAS DKO cells were not able to establish a growing culture in CMM_gly medium and the SCoAS RNAi induced cells showed a strong growth retardation followed by a significant decrease of ΔΨm. Indeed, the glycerol-induced reduction of the cytosolic ATP levels may create conditions under which the AAC activity is no more sufficient to compensate for the absence of mitochondrial substrate level phosphorylation (Figure 12D). Alternatively, we cannot exclude that AAC is reversed and exploits the cytosolic ATP pool and therefore further depletes the cytosolic levels of ATP causing the growth defect.

In summary, we can conclude that bloodstream forms exhibit an amazing flexibility in terms of cellular bioenergetics, which enables the parasite to quickly adapt and survive various challenging environments of its mammalian host by responding to sudden changes in intracellular ATP levels while still maintaining viable levels of the ΔΨm.

## Material and methods

### Trypanosoma cultures

A. *T. brucei brucei* bloodstream Lister 427 form (BSF 427) and genetic derivatives thereof were used in this study. The long slender monomorphic bloodstream forms were cultured in HMI-11, Creek Minimal Medium (CMM) containing 10 mM glucose (CMM_glc) or 10 mM glycerol (CMM_gly) supplemented with 10% heat-inactivated fetal bovine serum (FBS) at 37°C in the presence of 5% CO_2._ The genetically modified parasites were cultivated in HMI-11 medium in the presence of appropriate antibiotics to maintain their genetic background (G418 in 2.5 µg/ml, hygromycin in 5 µg/ml, puromycin in 0.1 µg/ml, phleomycin in 2.5 µg/ml, and tetracycline in 1 µg/ml). When needed, the cells were transferred to CMM_glc or CMM_gly media and maintained for two weeks before any experiments were performed (except for the experiment shown in Figure 11A). The cells were always kept in a logarithmic growth phase and harvested at a density of 0.7-1.4 x 10^6^ cells/ml.

### Plasmids and generation of genetically modified cell lines

The AAC double knock-out (DKO) and SCoAS DKO were generated by two rounds of homologous recombination using gene knock-out (KO) cassettes conferring either neomycin (G418) or hygromycin resistance. The gene cassettes were derived from the pLEW13 and pLEW90 vectors, respectively [54]. To direct the allele replacement, the KO casettes were flanked by short sequences of either AAC (Tb92710.14820/-30/-40) or SCoAS subunit β (Tb927.10.7410) 5´and 3´ untranslated regions (UTR) that were identified with TritrypDB. The UTR fragments were amplified by PCR from BSF 427 genomic (g)DNA with 5´UTR_forward and reverse or 3´UTR_forward and reverse primers (Supplementary Table S5). The amplicons were then digested with Not I and MluI restrictions enzymes (5´UTRs) or XbaI and StuI (3´UTRs) before sequentially ligated into the pLEW13 plasmid that contains genes for neomycin-resistance and T7 RNA polymerase gene. The final pLEW13_AAC_5´/3´UTRs and pLEW13_SCoAS_5´/3´UTRs constructs were linearized with Not I and electroporated with human T cell nucleofector solution (AMAXA) into BSF 427 to generate a single KO cell line. The transfected cells were serially diluted after 16 hours of recovery and selected with 2.5 µg/ml G418. To generate the double knock-out, the hygromycin-resistance cassette containing the tetracycline repressor under 10 % T7 RNAP promoter was excised from the pLEW90 vector with XhoI and Stu I restriction enzymes and used to replace the neomycin-resistance cassette from the pLEW13_AAC_5´/3´UTRs and pLEW13_SCoAS_5´/3´UTRs construct pre-digested with XhoI and SwaI endonucleases. The resulting constructs were linearized with NotI and the cassette was transfected to verified AAC_ and SCoAS_ single knock-out cell line followed by selection using hygromycin (5 µg/ml). AAC DKO and SCoAS DKO were grown in the presence of 2.5 µg/ml G418 and 5 µg/ml hygromycin.

To downregulate expression of SCoAS, DNA fragment corresponding to 591 bp target sequence was amplified by PCR from BSF427 gDNA using gene forward and reverse primers (Supplementary Table S5) extended with BamHI and HindIII restriction sites. The resulting PCR product was digested with the corresponding enzymes and inserted to digested p2T7-177 plasmid [55]. The verified plasmid was linearized with NotI and transfected into the single marker BSF 427 cell line which bears cassettes for T7 RNAP and tetracycline repressor under neomycin-resistance marker allowing for inducible expression of dsRNA using tetracycline. SCoAS RNAi were kept in G418 and phleomycin again with induction of RNAi by tetracycline.

To generate constructs for the constitutive expression of luciferase targeted to either cytosol or mitochondrial matrix, the luciferase gene was amplified by PCR using gene specific forward and reverse primers. To ensure mitochondrial localization of the luciferase, the mtLuc_FW primer was extended on its 5´termini with TbIscU mitochondrial targeting sequence [41]. The amplified *luc_mito* and *luc_cyto* genes were digested with BamHI and HindIII restriction enzymes and cloned into the modified pHD1344-tub-B5-3v5 vector (provided by J. Carnes and K. Stuart) that was pre-digested with the same enzymes to remove the original gene for TbKREPB5. The final plasmids pHD1344-tub-mtLUC-3v5 and pHD1344-tub-cytLUC-3v5 were linearized by NotI and transfected to BSF 427, AAC DKO and SCoAS DKO cell lines. The integration into the tubulin locus ensures constitutive expression of the gene of interest. Luciferase cell line in the BSF 427 background was grown in puromycin and in the background of AAC DKO and SCoAS DKO cells were grown in G418, hygromycin, and puromycin.

The AAC DKO/ASCT RNAi cell line was generated by transfecting Not1-linearized pLew-ASCT-SAS construct containing N-terminal fragment of *asct* gene [56] to AAC DKO cell line. AAC DKO/ASCT RNAi were grown in G418, hygromycin, and phleomycin with the induction of RNAi by tetracycline.

AAC DKO and SCoAS DKO addback cell lines were generated in the background of the respective DKO cells. The coding sequences of AAC and SCoAS were amplified from BSF 427 gDNA using specific forward and reverse primers that were extended with HindIII and BamHI restrictions sites. The amplified PCR products were digested, cloned to pT7_3v5 plasmid containing a gene for puromycin selection, linearized with NotI and transfected into the AAC or SCoAS DKO cells. Addback cell lines were grown in the presence of G418, hygromycin, and puromycin, and the expression of the ectopic alleles was initiated by the addition of 1µg/ml tetracycline.

SCoAS conditional DKO (cDKO) cell line was generated using SCoAS single KO cell line which was transfected with pT7_3v5_SCoAS linearized plasmid. After successful selection with puromycin, the second allele was replaced using the pLEW90_SCoAS_5´/ 3´UTRs construct. The transfection and selection was done in the presence of tetracycline ensuring expression of regulatable SCoAS. SCoAS cDKO was grown in the presence of tetracycline, G418, hygromycin, and phleomycin. Suppression of the ectopic allele expression was done by washing the cells twice in tetracycline-free media.

### Measurement of **ΔΨm** using flow cytometry

The ΔΨm was determined utilizing the red-fluorescent dye tetramethylrhodamine ethyl ester (TMRE, Invitrogen). Cells were grown in log-phase for a few days prior the experiment. In a specific case, the cells were pre-treated with oligomycin with the sublethal concentration of 250 µg/ml for 24 hours before the experiment. Then, in total, 5 × 10^6^ of oligomycin treated or untreated cells were pelleted (1,300 g, 10 min, room temperature), resuspended in 1 ml of the appropriate medium, incubated with 60 nM TMRE for 30 min at 37 °C, washed in PBS, resuspended in PBS-G (PBS, 6 mM glucose) and immediately analyzed by flow cytometry (BD FACS Canto II Instrument). In the case of oligomycin-treated cells, the 250 µg/ml of oligomycin was maintained in all buffers and washes. For each sample, 10,000 fluorescent events were collected. Treatment with the protonophore FCCP (20 μM) for 10 min was used as a control for mitochondrial membrane depolarization. Data were evaluated using BD FACSDiva (BD Company) software.

### SDS PAGE, Western blots, antibody production

Cell cultures were harvested at 1300g at 4°C for 10 minutes, washed with 1x PBS and the lysates were prepared at concentration 1×10^7^ cells/30 µl using 1xPBS, 6% sodium dodecyl sulfate, 300mM DTT, 150mM Tris HCl, 30% glycerol, and 0.02% Bromophenol Blue. Samples were boiled for 7 minutes at 97°C and stored at −20°C. Proteins were resolved on SDS-PAGE gels (BioRad 4568093, Invitrogen XP04202BOX) using 1×10^7^ cells/sample. Proteins were blotted onto PVDF membrane (Thermo Scientific) and probed with corresponding monoclonal (mAb) or polyclonal (pAb) antibodies. This was followed by probing with secondary HRP conjugated anti-mouse or anti-rabbit antibody (1:2000 dilution, SIGMA). Proteins were visualized using the Clarity Western ECL substrate (Bio-Rad 1705060EM) on a ChemiDoc instrument (Bio-Rad). The PageRuler pre-stained protein standard (Fermentas) was used to determine the size of the detected bands. AAC and SCoAS pAb were prepared for the purpose of this study. Open reading frames of AAC and SCoAS beta subunit were cloned in *E. Coli* expression plasmid pSKB3. Proteins were overexpressed in *E. Coli* BL21 cells, solubilized by sarkosyl, and purified by high-performance liquid chromatography. Antigens were sent to David’s Biotechnologie (Germany) for pAb production. Primary antibodies used in this paper are following: pAb anti-AAC (1:1000, 34kDa), pAb anti-SCoAS (1:1000, 45kDa), pAb anti-APRT (1:500, 26kDa), pAb anti-p18 (1:1000, 18kDa) and mAb anti-HSP70 (1:5000, 72kDa).

### Digitonin subcellular fractionation

Whole cell lysates (WCL) were prepared from BSF 427 for SCoAS localization and cell lines expressing mitochondrial (mito) or cytosolic (cyto) luciferase. For the digitonin fractionation, 1×10^8^ cells were harvested and washed with 1x PBS-G. Pellet was resuspended in 500 µl of SoTe (0.6 M Sorbitol, 2 mM EDTA, 20 mM Tris-HCl pH 7.5) and lysed with 500 µl of SoTe with 0.03% digitonin. Samples were incubated on ice for 5 minutes and centrifuged at 7000rpm for 3 minutes at 4°C. The supernatant was harvested as a cytosolic fraction and the pellet was resuspended in 1xPBS as a mitochondrial fraction. WCL and the fractions were resolved by SDS-PAGE.

### Measurement of **ΔΨm** using Safranin O dye

This method was performed as described previously [21]. Briefly, the in situ ΔΨm was measured using Safranin O dye (Sigma, S2255-25G). 2×10^7^ cells were centrifuged for 10 minutes at room temperature at 1300g and washed with ANT buffer containing 8 mM KCl, 110 mM K-gluconate, 10 mM Mannitol, 10 mM NaCl, 10 mM free acid HEPES, 10 mM K_2_HPO_4_, 0.015 mM EGTA potassium salt, 0.5 mg/ml fatty acid free bovine serum albumin, and 1.5 mM MgCl_2_ at pH 7.25. The cell pellet was resuspended with ANT buffer with 4 µM digitonin and 5 µM Safranin O. Fluorescence was recorded in a Hitachi F-7100 spectrofluorometer (Hitachi High Technologies) at a 5-Hz acquisition rate, using 495 nm excitation and 585 nm emission wavelengths. Samples were measured at room temperature and stirred during the experiment. Where indicated, 1 mM ATP as a substrate and inhibitors (1 µM CATR or 10 µM oligomycin) were added. Finally, the SF6847 uncoupler was used as a control of the maximal depolarization.

### SCoAS activity assay

The organellar pellet from 5×10^8^ digitonin-fractionated cells was resuspended in ANT buffer, sonicated 3 times for 10 sec at 20% power. The sample was spun down at 16,000 g for 5 min and supernatant containing mitochondrial matrix was subjected to SCoAS activity assay. The activity was assayed in ANT buffer in the presence of succinyl-CoA (0.2 mM), ADP (2 mM), Ellmańs reagent (5,5’-dithio-bis-[2-nitrobenzoic acid], DNTB (0.2 mM)) at 30°C. The released CoA-SH reacted with DNTB forming thio-nitrobenzoate anion (TNB) which production in time was monitored spectrophotometrically at 412 nM using a Tecan Infinite M200 plate reader. One unit is defined as an enzyme activity that converts one nanomole of succinyl CoA to CoA-SH in 1 minute per 1 mg of total protein.

### Alamar blue-based cell viability assay

Corresponding *T. brucei* cell lines were plated in transparent 96-well plates in a concentration of 5×10^3^ cells/ml in 200ul per well. Cells were grown in the presence of different CATR concentrations between 1µM to 500 µM or in the presence of TPMP between 0.3 nM to 500 µM for 72 hours in standard cultivation conditions. After 72 hours 20 µl of 125ug/ml of resazurin (Sigma, R7017-1G) was added to each well. After 24 hours the fluorescence was measured using Tecan Spark set up for 544 nm wavelength for excitation and 590 nm for emission. Data were analyzed using GraphPad Prism 9 to establish the EC_50_ values.

### In vivo ATP measurements

5×10^6^ cells with constitutively expressed luciferase in cytosol or mitochondrion were centrifuged at 1300g for 7 minutes at room temperature. Cells were washed with 1x PBS and resuspended in 160ul of HEPES-LUC buffer containing 20 mM HEPES, 116 mM NaCl, 5.6 mM KCl, 8 mM MgSO_4_ and 1.8 mM CaCl_2_ at pH 7.4. Cells were immediately placed in white bioluminescence 96 well plates, the background luminescence was measured by the Tecan Spark and 40 µl of 250 µM luciferin was injected in each sample. The luminescence was measured for 20 cycles and where indicated 10 mM glucose was injected and changes of luminescence were recorded for another 35 cycles.

### Animal experiments

Groups of 7 mice were used for each of the cell lines. Mice were infected by 1×10^5^ cells via 100ul intraperitoneal injection of either BSF 427, AAC DKO, AAC DKO addback, SCoAS DKO, and SCoAS DKO addback. Mice injected with tetracycline induced addback cell lines were put on doxycycline (200 µg/ml doxycycline and 5% sucrose) drinking regime 24 hours before injection. Blood samples from a tail prick were diluted in 1x SSC and 3.7% formaldehyde, and the parasitemia levels were counted using a hemocytometer (Counting Chamber CE NeubauerIMP DL). Parasitemia counts were observed for 15 days and mice displaying impaired health or a parasite load over 1×10^8^ cells/ml of blood were euthanized.

#### Exometabolomics

BSF 427, AAC DKO and SCoAS DKO trypanosomes were grown in log-phase in HMI11 media supplemented with the respective drugs. 1×10^7^ cells were collected by centrifugation at 1400 x g for 10 min at RT and washed with incubation buffer (PBS buffer supplemented with 5 g/L NaHCO_3_, pH 7.4) with the addition of 1mM of the respective carbon source. Next, the cells were incubated in preheated plates until the cells manage to keep cell integrity (validated by microscopic observation, appr. 2.5 hours) at 37^0^C with incubation buffer containing uniformly labled [U-13C]-glucose (4 mM) in the presence or absence of the 4 mM amino acid threonine in a total volume of 1ml. The same experiment was carried out with ordinary ^12^C glucose as the only carbon source. Following centrifugation at 8000xg for 1 min at RT, the supernatant was collected and a proton NMR (^1^H-NMR) spectra analysis was performed as described in [31].

### LC-MS Endometabolomics

Samples were prepared as described previously (3). 5×10^7^ cells for each sample were rapidly cooled down in an ethanol-dry ice bath, centrifuged at 1300g for 10 minutes at 4°C, and washed with 1x PBS. Pellet was resuspended in 100ul of extraction solvent containing chloroform, methanol, and water (1:3:1 volume ratio). Samples were shaken for 1 hour at 4°C, pelleted at 13000g for 10 minutes at 4°C and the supernatants were stored at −80°C until analysis. The used metabolomic methods were described in detail elsewhere [57, 58]. Briefly, an Orbitrap Q Exactive Plus mass spectrometer coupled to an LC Dionex Ultimate 3000 (all Thermo Fisher Scientific, San Jose, CA, USA) was used for metabolite profiling. LC condition: column SeQuant ZIC-pHILIC 150mm x 4.6 mm i.d., 5μm, (Merck KGaA, Darmstadt, Germany); flow rate of 450 μL/min; injection volume of 5 μL; column temperature of 35°C; mobile phase A = acetonitrile and B = 20 mmol/L aqueous ammonium carbonate (pH = 9.2; adjusted with NH_4_OH); gradient: 0 min, 20% B; 20 min, 80% B; 20.1 min, 95% B; 23.3 min, 95% B; 23.4 min, 20% B; 30.0 min 20% B. The Q-Exactive settings were: mass range 70-1050 Daltons; 70 000 resolution; electrospray ion source operated in the positive and negative modes. Data were processed using Xcalibur™ software, version 4.0 (Thermo Fisher Scientific, San Jose, CA, USA), and an in-house developed Metabolite Mapper® platform containing more than 1500 metabolites mannually annotated against authentic standards.

### Mass spectrometry sample preparation, MS measurement, and proteomics data analysis

A. *T*. *brucei* BSF 427, SCoAS and AAC DKO cells (1×10^8^ cells/replicate) were washed three times in 10 mL of phosphate-buffered saline (PBS) and lysed in 6% sodium dodecyl sulfate (SDS), 300 mM DTT, and150 mM Tris-HCl (pH 6.8), 30% glycerol, and 0.02% Bromophenol Blue. Samples were loadedon a NOVEX NuPage 4%–12% gradient gel (Thermo Fisher Scientific, Waltham, MA), run for10 minutes at 180 V, and stained with Coommassie G250. Each lane was cut and the mincedgel pieces were transferred to an Eppendorf tube for destaining with 50% ethanol/50 mM ABC buffer pH 8.0. The gel pieces were dried and subsequently reduced (10 mM DTT/50 mM ABC buffer pH 8.0), alkylated (55 mM iodoacetamide/50 mM ABC buffer pH 8.0), and digested with 1 μg trypsin overnight at 37˚C. The tryptic peptides were eluted from the gel pieces with pure acetonitrile and stored on a StageTip. The proteomic measurement was performed on a Q Exactive Plus mass spectrometer (Thermo Fisher Scientific, Waltham, MA) with an online-mounted C18-packed capillary column (New Objective, Woburn, MA) by eluting along a 225-minute gradient of 2% to 40% acetonitrile using an EasyLC 1000 uHPLC system (Thermo Fisher Scientific, Waltham, MA). The mass spectrometer was operated with a top10 data-dependent acquisition (DDA) mode. Data analysis was performed in MaxQuant version 1.5.2.8 using the tritrypDB- 43_TbruceiLister427_2018_AnnotatedProteins database (16,869 entries) and standard settings, except activating the match between run feature and the label-free quantification (LFQ) algorithm. Protein groups marked as contaminants, reverse entries, and only identified by site were removed prior to bioinformatics analysis, as well as protein groups with less than 2 peptides (minimum 1 unique). Additional information like gene names and descriptions were extracted from the fasta header and attached to the individual protein groups.

### Statistical analysis

The number of replicates, controls, and statistical tests are in accordance with published studies employing comparable techniques and are generally accepted in the field. Statistical differences were analyzed with Prism software (version 8.2.1, GraphPad software). Comparisons of two groups were calculated with two-tailed paired *t* test. A *P* value of less than 0.05 was considered statistically significant. Quantitative mass spectrometry experiments were performed in four biological replicates.

## Supporting information

**Table S1.** Proteomic analysis of AAC DKO and SCoAS DKO cells. Sheet 1 contains Tb927 gene IDs and description for 3654 protein groups identified by a minimum of 2 peptides (1 unique) and present in at least two out of four replications. Sheet 2 contains protein groups identified in BSF 427 cells and compared to AAC DKO. Sheet 3 contains protein groups differentially expressed (log2 fold change < - 0.4, log2 fold change > 0.5) which passed threshold of *p-*value of 0.05. Sheet 4 contains protein groups identified in BSF 427 cells and compared to SCoAS DKO. Sheet 5 contains protein groups differentially expressed (log2 fold change < - 0.4, log2 fold change > 0.5) which passed threshold of *p-*value of 0.05.

**Table S2.** Metabolomic analysis of AAC DKO cells. LC-MS metabolomic data.

**Table S3.** Metabolomic analysis of AAC DKO cells. LC-MS metabolomic data.

**Table S4.** Excreted end-products from metabolism of glucose and threonine in the BSF trypanosomes. Parasites were incubated with 4 mM glucose or with [U-_13_C]-glucose with or without 4 mM threonine. ICS (internal carbon source): intracellular carbon source of unknown origin metabolized by the BSF trypanosomes. Amounts of end-products excreted (here malate) from the carbon source indicated in brackets, expressed as nmoles excreted per h and per 108 cells. *nd*: not detectable.

**Table S5.** List of oligonucleotides used in the study.

## Funding

This work was supported by Czech Science Foundation grant 20-14409S, by European Regional Development Fund (ERDF) and Ministry of Education, Youth and Sport (MEYS) project CZ.02.1.01/0.0/0.0/16_019/0000759 and by the European Research Council (ERC) under the European Union’s Horizon 2020 Research and Innovation Program (grant agreement no. 101044951) to AZ. FB and EP were supported by the Centre National de la Recherche Scientifique (CNRS), the Université de Bordeaux, the Agence Nationale de la Recherche (ANR) through grants ADIPOTRYP (ANR19-CE15-0004-01) of the ANR "Générique" call, the Laboratoire d’Excellence (LabEx) ParaFrap ANR-11-LABX-0024 and the "Fondation pour le Recherche Médicale" (FRM) ("Equipe FRM", grant n°EQU201903007845).

## Author contributions

## Notes

### Competing Interest Statement

The authors have declared no competing interest.

